# Symmetry transitions during gating of the TRPV2 ion channel in lipid membranes

**DOI:** 10.1101/545293

**Authors:** Lejla Zubcevic, Allen L. Hsu, Mario J. Borgnia, Seok-Yong Lee

**Author notes:** Correspondence to: S.-Y. Lee.

## Abstract

The Transient Receptor Potential Vanilloid 2 (TRPV2) channel is a member of the temperature-sensing thermoTRPV family. Recent advances in cryo-electronmicroscopy (cryo-EM) and X-ray crystallography have provided many important insights into the gating mechanisms of thermoTRPV channels. Interestingly, crystallographic studies of ligand-dependent TRPV2 gating have shown that the TRPV2 channel adopts two-fold symmetric arrangements during the gating cycle. However, it was unclear if crystal packing forces played a role in stabilizing the two-fold symmetric arrangement of the channel. Here we employ cryo-EM to elucidate the structure of full-length rabbit TRPV2 in complex with the agonist resiniferatoxin (RTx) in nanodiscs and amphipol. We show that RTx induces two-fold symmetric conformations of TRPV2 in both environments. However, the two-fold symmetry is more pronounced in the native-like lipid environment of the nanodiscs. Our data offers insights into a gating pathway in TRPV2 involving symmetry transitions.

## Introduction

Transient Receptor Potential V (TRPV) channels are part of the larger TRP channel family which play important roles in numerous physiological processes^1^. A subset of TRPV channels, including subtypes TRPV1-TRPV4, possess an intrinsic capability to sense heat and are therefore referred to as thermoTRPV channels ^2–5^. TRPV1-TRPV4 are non-selective cation channels which play important physiological roles in sensing noxious heat ^6–9^, maintaining cardiac structure ^10^ and maintaining skin ^11–13^, hair ^14–16^ and bone physiology ^17^. A distinctive feature of TRPV1 and TRPV2 is their permeability to large organic cations^18^, such as the cationic dye YO-PRO-1 and the sodium channel blocker QX-314. This feature has led to proposals to utilize these channels as conduits for delivering small molecules to intracellular targets^19^. TRPV1 and TRPV2 possess two activation gates, one at the selectivity filter (termed the SF gate) and second one at the intracellular mouth of the pore (termed the common gate)^20–22^. Both gates must open widely to accommodate the passage of large organic cations. However, the mechanism that enables such opening was long unclear. In order to study the permeation of both metal and large organic cations in TRPV2, we recently crystallized the rabbit resiniferatoxin (RTx)-sensitive^23^ TRPV2 channel with a truncation in the pore turret in the presence of the agonist RTx^24^. This study led to the revelation that the binding of RTx leads to a two-fold symmetric (C2) opening at the selectivity filter gate that is wide enough to permeate YO-PRO-1. This unexpected result offered the first experimental evidence that the homotetrameric TRPV2 can adopt C2 symmetric conformations during the gating cycle. However, it was unclear if crystal contacts or the crystallization conditions (e.g. high concentration of Ca^2+^) played a role in stabilizing the C2 symmetry. In addition, the minimal TRPV2 construct used in the crystallographic study lacked the pore turret, a region that is not essential for function^20,21,24,25^, but had previously been shown to have a modulatory effect on gating in TRPV1 and TRPV2^26,27^. It was uncertain if the absence of this region in our crystallographic study affected the symmetry of the channel.

In order to answer these questions and further study the role of two-fold symmetry in TRPV channel gating, we conducted cryo-electronmicroscopy (cryo-EM) studies of the full-length, RTx-sensitive rabbit TRPV2^23^ channel reconstituted into nanodiscs and amphipol. We present three structures of the TRPV2/RTx complex, one obtained in nanodiscs (TRPV2_RTx-ND_) and three in amphipol (TRPV2_RTx-APOL 1-3_) determined to 3.8 Å, 2.9 Å, 3.3 Å and 4.2 Å resolution, respectively. Our data shows that binding of RTx induces C2 symmetric conformations in TRPV2, but the degree of symmetry reduction depends on the environment in which the channel is reconstituted. C2 symmetry is particularly pronounced in the dataset collected from nanodisc-reconstituted TRPV2, which better approximates the physiological environment of the channel. Moreover, the data offers further insights into the allosteric coupling between the RTx binding site and the activation gates in TRPV2, confirms the critical role of the S4-S5 linker π-helix (S4-S5_π-hinge_) in ligand-dependent gating of TRPV2, and provides a glimpse of the conformational landscape of TRPV2 gating.

## Results

In order to capture the RTx-induced gating transitions in the rabbit TRPV2 channel, we conducted cryo-EM studies of the TRPV2/RTx complex reconstituted into amphipol (TRPV2_RTx-APOL_) and nanodiscs (TRPV2_RTx-ND._) Amphipols 28 have been a useful tool in structural studies of membrane proteins, and especially TRP channels^20,21,29–33^. Indeed, Amphipol A8-35 enabled the very first structural determination of the TRPV2 channel^21^. Nanodiscs, on the other hand, represent the closest *in vitro* approximation to the native lipid membranes used in structural studies^34^. The data was processed using Relion^35^ (Methods), with no symmetry imposed during classification and 3D reconstruction of the particles in order to avoid obscuring any classes with lower symmetry (C1-C2) that might exist in the sample. Symmetry was only imposed in the last step of the refinement and only if the 3D reconstructions showed clear two-fold (C2) or four-fold (C4) symmetry (Figure Supplements 1–2). Classification of the TRPV2_RTx-APOL_ sample revealed the presence three different conformations: one C4 symmetric and two distinct C2 symmetric classes refined to 2.9 Å, 3.3 Å and 4.2 Å, respectively (Figure 1, Figure Supplement 1). By contrast, 3D classification of the TRPV2_RTx-ND_ converged on a single C2 symmetric conformation resolved to 3.8 Å (Figure 1, Figure Supplement 2). All four maps were of sufficient quality to enable placement of individual structural motifs with confidence (Figure Supplements 3-6) and the models for all four structures were built to good overall geometry (Table 1).

**Table 1.**
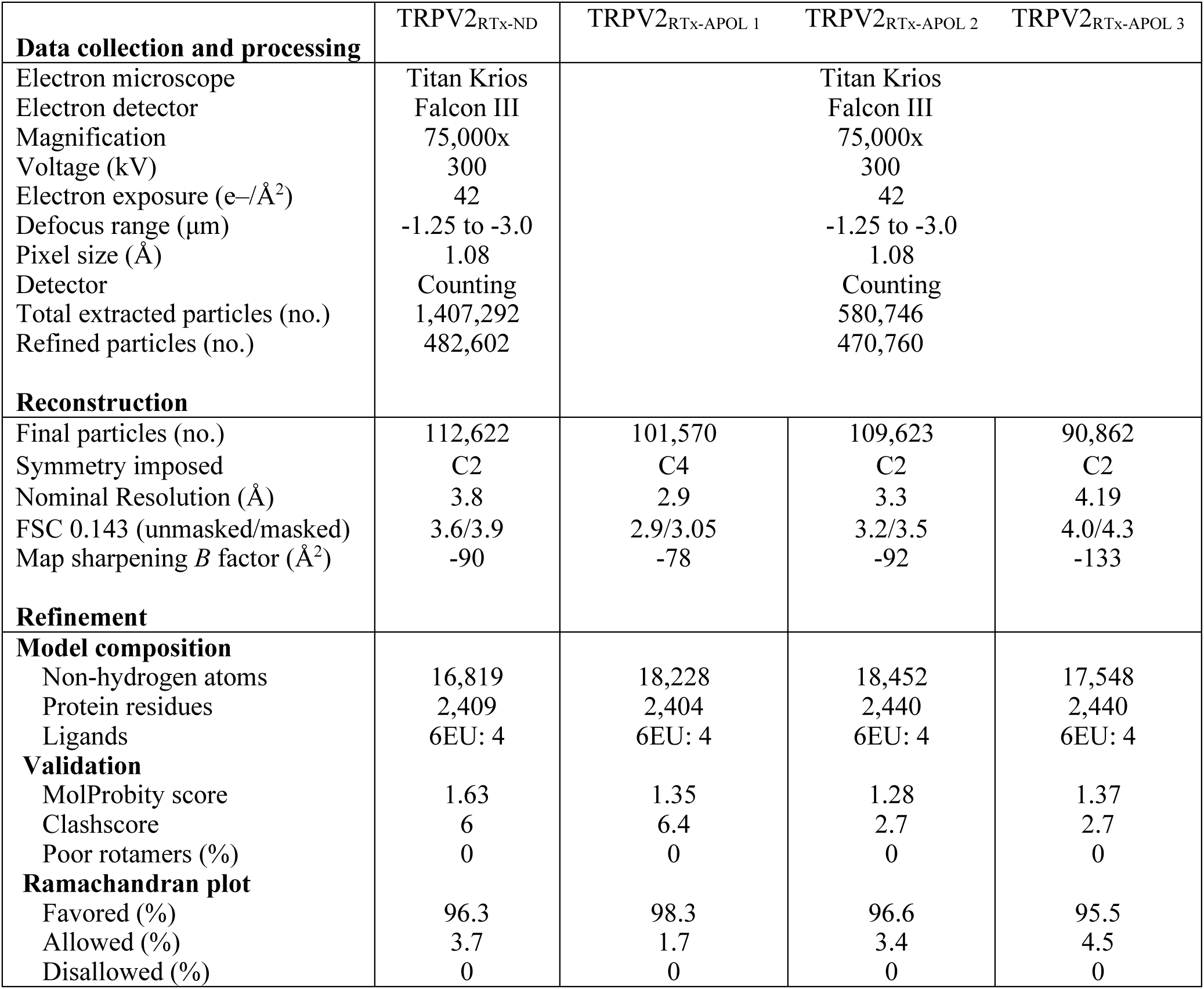
Data collection and refinement statistics

**Figure 1.**
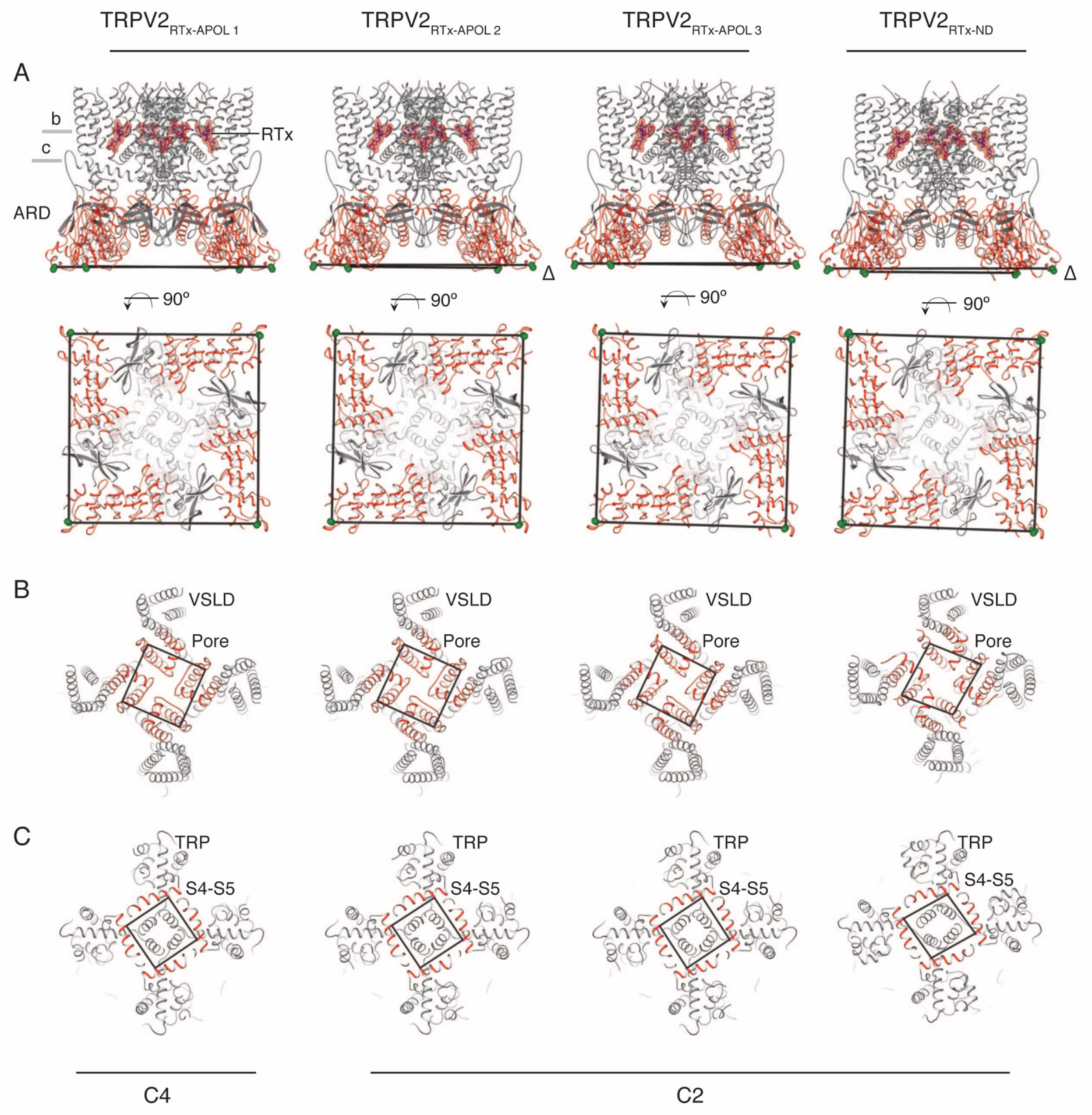
Overview of TRPV2_RTx-APOL_ and TRPV2_RTx-ND_ structures. **A,** Orthogonal view of TRPV2_RTx-APOL_ 1-3 and TRPV2_RTx-ND_ structures. TM domains are colored in gray and the cytoplasmic domains (ARD and C-terminal domain) are colored in red. RTx is shown in stick and sphere representation and colored in red. Lines drawn between diagonally opposite ARDs (residue E95, shown in green spheres) illustrate the relative position of ARDs in the tetramer. **B,** Bottom-up view of the ARD (red). Lines drawn between residues E95 (green spheres) illustrate the symmetry and rotation of the ARD assemblies. **C,** Top view of the channel (red). Lines drawn between residues V620 in the S6 helix illustrate the symmetry within the pore domain (red). **D,** Lines drawn between residues Y523 show symmetry in the S4-S5 linker (red).

### The transmembrane domains of TRPV2_RTx-APOL_ are trapped in a closed conformation

Unexpectedly, the transmembrane domains (TM) of the three structures obtained from amphipol-reconstituted TRPV2, TRPV2_RTx-APOL_ 1-3, show similarity to our previously solved cryo-EM structure of TRPV2 in its apo form^21^ (TRPV2_APO_) and adopt non-conducting conformations (Figure Supplement 7). While fully bound to RTx, the TM domains of TRPV2_RTx-APOL_ 1 and TRPV2_RTx-APOL_ 2 structures largely retain C4 symmetry (Figure 1 and Figure Supplement 8). However, the TMs of TRPV2_RTx-APOL_ 3 exhibit a slight departure from C4 symmetry in the pore (Figure Supplement 9). The effects of RTx on the TRPV2_RTx-APOL_ are particularly obvious in the ankyrin repeat domains (ARD) of the two-fold symmetric TRPV2_RTx-APOL_ 2 and TRPV2_RTx-APOL_ 3 which display pronounced broken symmetry and a range of rotational states (Figure 1, Figure Supplements 9-10).

In order to determine the effect of RTx on the TRPV2_RTx-APOL_ sample, we aligned TRPV2_RTx-APOL_ 1 with TRPV2_APO_. The transmembrane helices S1-S6 of the two channels aligned remarkably well (Cα R.M.S.D =0.86) (Figure Supplement 8). However, RTx binding induces a 5° clockwise rotation of the ARD when viewed from the extracellular space and a ~10 Å lateral widening of the cytoplasmic assembly (Figure Supplement 8). In addition, RTx causes a conformational change in the S4-S5 linker (Figure Supplement 8), as well as a displacement of the TRP domain (Figure Supplement 8). The conformational change in the S4-S5 linker is caused by the introduction of a π-helical turn at the junction of the S4-S5 linker and the S5 helix in the TRPV2_RTx-APOL_ 1 structure (S4-S5_π-hinge_), which is absent in TRPV2_APO_ (Figure Supplement 8). This observation concurs with our previous finding that RTx binding elicits a conformational change in the S4-S5 linker, and that the S4-S5_π-hinge_ is critical for ligand-dependent gating in TRPV2^24^. In TRPV2_RTx-APOL_ 3, slight C2 symmetry is observed in the TM domains and is evident in the SF gate, PH and the S4-S5 linker (Figure Supplement 9).

Nevertheless, the RTx-induced conformational changes in the S4-S5 linker are not efficiently propagated to the TM in the TRPV2_RTx-APOL_ structures, and they fail to open either of the two activation gates (Figure Supplement 7). Instead, RTx only effects changes in its immediate binding site above the S4-S5 linker and in the parts of the channel not bound by amphipol, strongly suggesting that the polymer constricts the TM and prevents conformational changes at the S4-S5 linker and the ARD from propagating to the TM domain. The fact that the TRPV2/RTx complex is stabilized in multiple distinct closed states with different arrangements of the ARD assembly (Figure 1, Figure Supplements 9-10) suggests that the conformational changes in the ARD might represent low-energy, pre-open states that can be achieved without substantial changes in the TM domains.

Interestingly, metal ions are not visualized in the pores of any of the TRPV2_RTx-APOL_ structures, despite the high resolutions obtained in this study. Whether this is the result of cryo-EM experimental conditions is unclear, but thus far metal ions occupying the SF and the pores of thermoTRPV channels have only been captured in structures obtained by X-ray crystallography^24^.

### RTx induces a break in symmetry in TRPV2_RTx-ND_

In stark contrast to the amphipol-reconstituted channel, reconstitution in nanodiscs revealed that RTx binding induces widespread C2 symmetry in TRPV2 which extends throughout the channel. Both activation gates in TRPV2_RTx-ND_ adopt C2 symmetric arrangements (Figure 2a). The pore helices of the SF gate are arranged so that the carbonyl oxygens of the selectivity filter in subunits B and D line the entry to the pore while pore helices of subunits A and C are tilted away from the permeation pathway. This arrangement creates a large C2 symmetric opening where the narrowest constriction between SF gate residues in diametrically opposing subunits A and C and B and D is ~ 11 Å and ~8.3 Å, respectively. This results in an SF gate with ample room to accommodate large organic cations (Figure 2b). A closer look at the pore helices reveals that this arrangement in the SF gate is achieved through a ~27° swivel of the subunit A pore helix, which brings the N-terminal part of the helix closer to S5 while distancing it from S6 (Figure 2c). The position of the pore helices controls the size and the shape of the SF gate, thereby exerting dynamic control over ion permeation in TRPV2. While the SF gate is widely open, the conformation of the common gate is a hybrid of closed and open states. In subunits A and C, the S6 helix adopts an α-helical, closed conformation, while a secondary structure transition in S6 of subunits B and D results in the presence of a π-helical turn which bends the helix and opens the common gate (Figure 2a).

**Figure 2.**
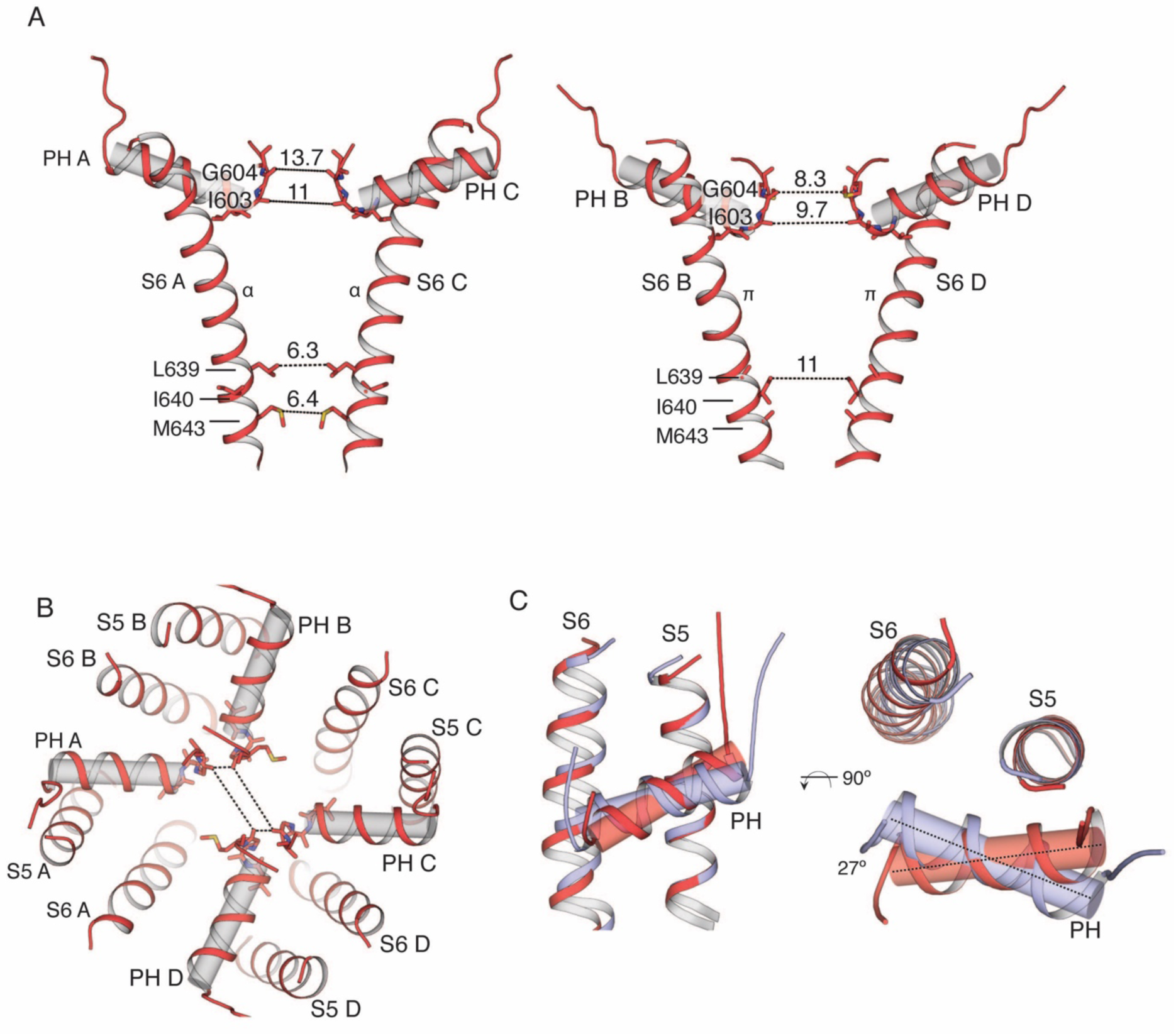
Overview of the pore in the TRPV2_RTx-ND_ structure. **A,** S6 and pore helices of subunits A and C (left) and subunits B and D (right). Pore helices are shown in both cartoon and cylinder representation (gray). Dashed lines and values represent distances between the indicated residues. S6 helices in A and C are α-helical, while a π-helical turn is introduced in subunits B and D. **B,** Top view of the TRPV2_RTx-ND_ pore, with pore helices shown in both cartoon and cylinder representation. Dashed lines illustrate the distances between residues G604 in the selectivity filter. **c,** Overlay of the TRPV2_RTx-ND_ pore domains (S5, S6 and pore helices). Subunit A is shown in red and subunit B in violet. The pore helix of subunit A swivels by ~27° relative to subunit B.

In order to establish the origin of the C2 symmetry in the TRPV2_RTx-ND_ structure, we aligned subunits A and B (Cα R.M.S.D =0.96) (Figure Supplement 11). Similar to our previous findings, this alignment shows that the two subunits diverge at the S4-S5 linker and the PH and indicates that rotation of subunits around the S4-S5_π-hinge_ appears to result in the distinct C2 symmetric arrangement observed in TRPV2_RTx-ND_ (Figure Supplement 11).

When compared to the TRPV2_APO_, the TM domains of the TRPV2_RTx-ND_ structure appear to contract in an asymmetric manner (Figure 3a), while the ARD assembly expands by ~10 Å and rotates by 3° (Figure 3b). The TM domains and the ARDs appear to move as a single rigid body, which is evident when individual subunits from TRPV2_APO_ and TRPV2_RTx-ND_ are superposed (Cα R.M.S.D =1.9 Å) to reveal that only the S4-S5 linker and the pore helix deviate significantly in the two structures (Figure 3c). This coupled movement of the TM and ARD indicates that RTx-binding to TRPV2 in lipid membranes induces a rigid-body rotation of the entire subunit that originates at the S4-S5_π-hinge_ (Figure 3d-e).

**Figure 3.**
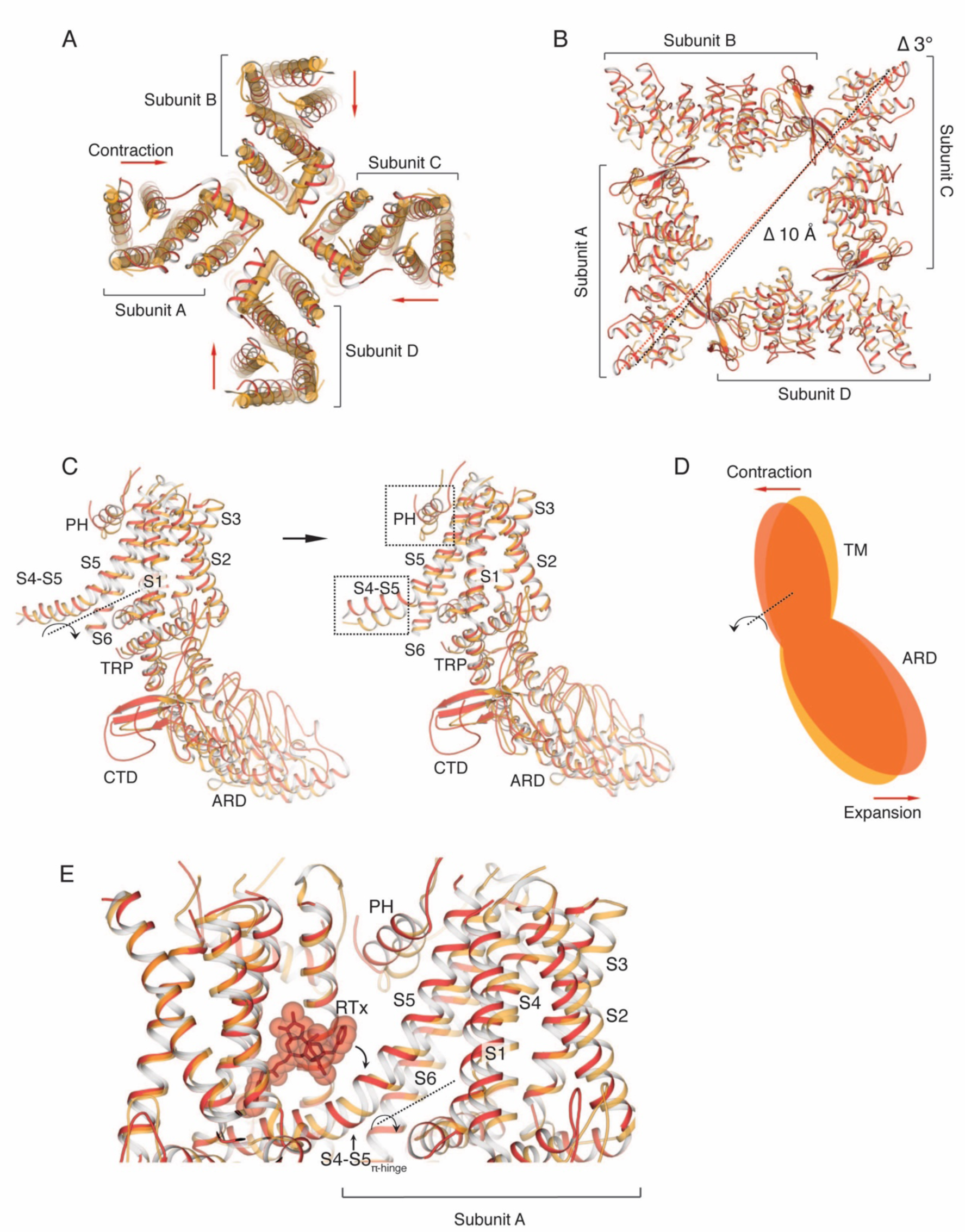
Comparison of TRPV2_RTx-ND_ (red) and TRPV2_APO_ (orange). **A,** Overlay of TRPV2_RTx-ND_ and TRPV2_APO_, top view. TRPV2_RTx-ND_ is shown in cartoon representation and TRPV2_APO_ as cylinders. Relative to TRPV2_APO_, the TM subunits of TRPV2_RTx-ND_ exhibit contraction (red arrows). **B,** Top view of the ARDs in TRPV2_RTx-ND_ and TRPV2_APO_. TM helices are removed for ease of viewing. Dashed lines represent distances between residues T100, showing a 10 Å expansion and 3° rotation of the TRPV2_RTx-ND_ ARD assembly relative to TRPV2_APO_. **C,** A rigid-body rotation of TRPV2_RTx-ND_ subunit B around the S4-S5 linker achieves alignment with the subunit B from TRPV2_APO_. Following alignment, only the S4-S5 linkers and the pore helices (PH) diverge in the two subunits (dashed box). **D,** Cartoon illustrating how the movements of the TM and the ARD in TRPV2_RTx-ND_ are coupled. The red and orange shapes represent a single subunit of TRPV2_RTx-ND_ and TRPV2_APO,_ respectively. The rotation of the subunit is manifested as “contraction” in the TM domains “expansion” of the ARD. **E,** RTx binding in the vanilloid binding pocket exerts force on the S4-S5 linker, changing the conformation of the junction from α- to π-helix, and induces the rotation of the subunit around the S4-S5__π-hinge__.

Interestingly, the TRPV2_RTx-ND_ structure exhibits different degrees of reduced symmetry from the previously determined crystal structure of TRPV2 in complex with RTx (TRPV2_RTx-XTAL_)^24^. Compared to the TRPV2_RTx-XTAL,_ the TM domains of subunits A and C in TRPV2_RTx-ND_ are widened, while those of subunits B and D exhibit a contraction (Figure 4a). This conformational change, which stems from rotation of individual TRPV2_RTx-ND_ subunits around the S4-S5_π-hinge_ (Figure Supplement 12), results in an overall fold that is closer to C4 symmetry than that of the TRPV2_RTx-XTAL_ (Figure 4b). However, while the TRPV2_RTx-ND_ helices S1-S6 adopt a more C4 symmetric arrangement, the pore helices and the SF gate remain distinctly C2 symmetric (Figure 4c). Remarkably, the SF gate of TRPV2_RTx-ND_ is wider than in TRPV2_RTx-XTAL,_ and the two structures display different C2 symmetric openings at the SF gate (Figure 4c). The two different conformations result from both the different arrangements of subunits and changes in the position and tilt angle of the pore helices (Figure 4d-e). In the TRPV2_RTx-XTAL_ structure, the pore helices of subunits B and D, which assume a widened conformation, are free of interactions with the pore domain, while a network of hydrogen bonds (Y542-T602-Y627) in subunits A and C tethers the pore helices to S5 and S6. Our previous work showed that disruption of these hydrogen bonds is detrimental to the permeation of large organic cations, but has no effect on permeation of metal ions^24^. Interestingly, the hydrogen bond triad is disrupted in all four subunits of the TRPV2_RTx-ND_ structure (Figure Supplement 13). Nevertheless, the SF gate assumes a fully open state that can easily accommodate passage of a large cation. This suggests that the hydrogen bond triad, while not a feature of the fully open SF gate, is an essential part of the transition between closed and open states of the channel.

**Figure 4.**
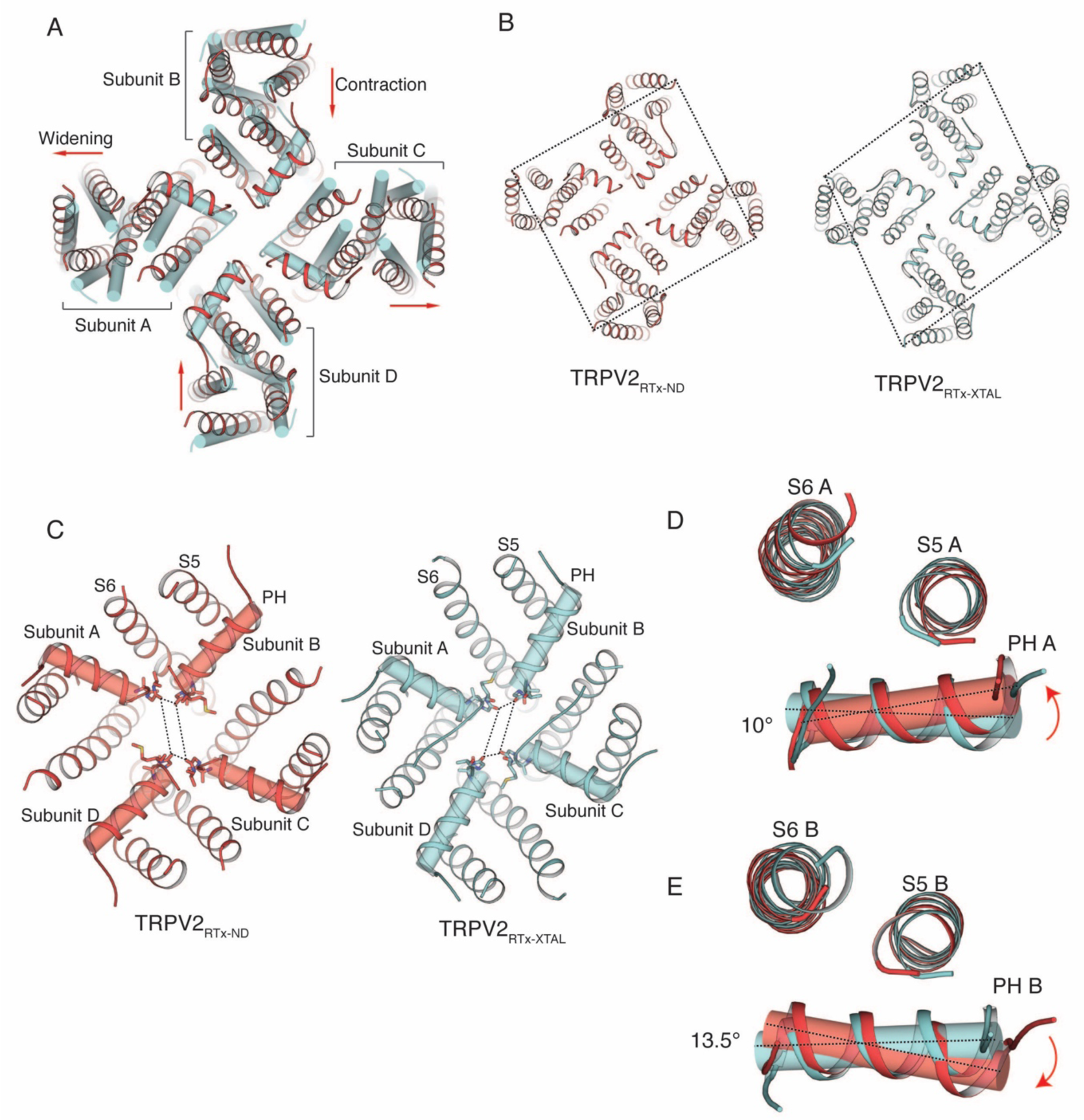
Comparison of TRPV2_RTx-ND_ (red) and TRPV2_RTx-XTAL_ (cyan). **A,** Overlay of TRPV2_RTx-ND_ and TRPV2_RTx-XTAL,_ top view. TRPV2_RTx-ND_ is shown in cartoon representation and TRPV2_RTx-XTAL_ as cylinders. Relative to TRPV2_RTx-APOL_ 1, subunits A and C of TRPV2_RTx-ND_ are widened, while subunits B and D exhibit contraction (red arrows). **B,** Comparison of two-fold symmetry in TRPV2_RTx-ND_ and TRPV2_RTx-XTAL_. Dashed lines represent distances between residues A427. **C,** Top view of the SF gate in TRPV2_RTx-ND_ and TRPV2_RTx-XTAL_. Pore helices are shown in both cartoon and cylinder representation. Dashed lines represent distances between residues G604 in the selectivity filter. **D-E,** Overlay of the pore domains of TRPV2_RTx-ND_ and TRPV2_RTx-XTAL_ subunit A **(D)** and subunit B **(E)** show that the pore helices A and B in TRPV2_RTx-ND_ swivel by ~10° and 13.5°, respectively, compared to TRPV2_RTx-XTAL_.

Despite the use of a full-length rabbit TRPV2 construct in this study, we were not able to confidently resolve the entire loop connecting S5 to the pore helix known as the “pore turret”. Interestingly, a recent structure of rat TRPV2 with the pore turret resolved showed that this region, which contains a large number of charged and polar residues, occupies the space within the membrane plane between S5 and the Voltage Sensing Like Domain (VSLD)^27^. While the density in our cryo-EM maps was not of sufficient quality to build the entire pore turret with confidence, we do observe density following the S5 helix and preceding the pore helix. However, the direction of this density is perpendicular to the membrane and does not agree with the structure reported for rat TRPV2 (Figure Supplement 14). Indeed, the pore turret is amongst the least conserved regions amongst the TRPV2 orthologs, and the variations in its sequence might be reflective of different conformations in TRPV2 channels of different species. Nevertheless, our study clearly shows that the omission of this region from the construct used in the crystallographic study of the TRPV2/RTx complex is not the cause of the C2 symmetry.

While both TRPV2_RTx-ND_ and TRPV2_RTx-XTAL_ structures adopt C2 symmetry, the distinct arrangement of subunits within the two channels suggests that the structures represent different functional states. We propose that TRPV2_RTx-XTAL_ precedes TRPV2_RTx-ND_ in the conformational activation trajectory based on two observations. Firstly, the common gate is fully closed in the TRPV2_RTx-XTAL_ while it adopts a partially open state in TRPV2_RTx-ND_ (Figure Supplement 15). Secondly, our previous studies have shown that the hydrogen bond network between S5 and S6 and the pore helix is essential for the channel’s ability to transition to a fully open SF gate that can accommodate large organic cations^24^. Nevertheless, in TRPV2_RTx-ND_ the pore helices do not interact with S5 and S6 and the SF gate is fully open. Therefore, the conformational step that requires the presence of the hydrogen bond triad must precede the open SF gate conformation seen in TRPV2_RTx-ND_.

## Discussion

Here we have conducted a study that reveals symmetry transitions associated with gating of the TRPV2 channel by RTx. Interestingly, our data shows that RTx induces C2 symmetric conformations of TRPV2 in both amphipol and nanodiscs, and it thereby negates the hypothetical role of crystallization artefacts and crystal packing bias in stabilising two-fold symmetry. Similarly, C2 symmetry in TRPV2 is independent of the presence or absence of the pore turret region, suggesting that this region does not play an essential role in the regulation of the SF gate in rabbit TRPV2. Our study, similar to a previously published study of the magnesium channel CorA^36^, also emphasizes the notion that careful inspection of the intermediate maps and conservative application of symmetry during refinement of cryo-EM data can result in valuable insights into gating transitions and intermediate states. In addition, we have also investigated how amphipols and nanodiscs affect the conformational space that can be accessed during ligand gating of TRPV2.

While both TRPV2_RTx-APOL_ and TRPV2_RTx-ND_ are C2 symmetric, the two-fold symmetry in TRPV2_RTx-APOL_ is confined to regions that are not bound by the amphipol polymer. This is evident in the fact that the TM domains, which are in contact with the amphipol, largely retain four-fold symmetry and the two gates remain firmly closed, while the ARD exhibit symmetry breaking, rotation and lateral expansion. These data, while adding valuable data points to the conformational landscape of TRPV2, also illustrate the caveats of using amphipols in studies of conformational changes in the transmembrane domains of proteins, as they appear to constrict the TM domains and stabilize low-energy pre-open states. By contrast, the TRPV2_RTx-ND_ dataset yielded a single, two-fold symmetric structure thus giving strong evidence that RTx stabilizes two-fold symmetric conformational states in the TRPV2 channel in lipid membranes. The ARDs in the TRPV2_RTx-ND_ structure echo the conformational changes observed in TRPV2_RTx-APOL._ However, in nanodiscs TRPV2 is captured with its SF gate fully open and its common gate in a conformation that reflects a mixture of open and closed states. In this structure, the opening of the SF gate occurs according to a mechanism previously observed in the crystallographic study of the TRPV2/RTx complex where RTx binding in the vanilloid pocket, above the S4-S5_π-hinge,_ induces a rigid body rotation of the entire subunit. In turn, the rotation causes a break in the hydrogen bond network between the pore helix and helices S5 and S6, allowing the pore helices to reposition and the SF gate to open^24^.

Interestingly, however, the TRPV2_RTx-ND_ structure differs from the previously obtained TRPV2_RTx-XTAL_. While both structures assume C2 symmetric conformations, the TRPV2_RTx-ND_ channel appears to make a return towards C4 symmetry. Because the SF gate in TRPV2_RTx-ND_ is fully open, and two of its S6 helices contain a π-helix and adopt an open conformation, we reason that TRPV2_RTx-ND_ follows the TRPV2_RTx-XTAL_ structure in the conformational trajectory of the channel. Therefore, it is possible that TRPV2, as it travels towards the final open state where both the SF and the common gate are fully open, would adopt further conformations that increasingly approximate C4 symmetry (Figure 5). However, it is interesting to note that while the overall fold of TRPV2_RTx-ND_ indeed is more C4 symmetric than that of TRPV2_RTx-XTAL_, the extent of C2 symmetry is not diminished in its SF gate. Because the symmetry of the SF gate does not appear to be dictated by the symmetry of the overall channel, we cannot exclude the possibility that the final open state might indeed possess a C2 symmetric SF gate while otherwise adopting a nearly C4 symmetric conformation. Our previous functional studies have shown that C2 symmetric states are critical for the channel’s ability to conduct large organic cations and consequently for the full opening of the SF gate^24^. Hence, the channel might be utilizing C2 symmetric states as means to achieve full opening in a step-wise manner. Similar C2 symmetric states elicited by ligand binding have been observed in TRPV3^33^ and TRPM2^37^ channels, which opens up the possibility that C2 symmetry might be widely associated with gating in members of the TRP channel superfamily. Intriguingly, a recent cryo-EM study of the human BK channel reconstituted in liposomes showed that this channel also enters C2 symmetric states^38^, suggesting that two-fold symmetry might also play a role in the molecular mechanisms of other tetrameric ion channels.

**Figure 5.**
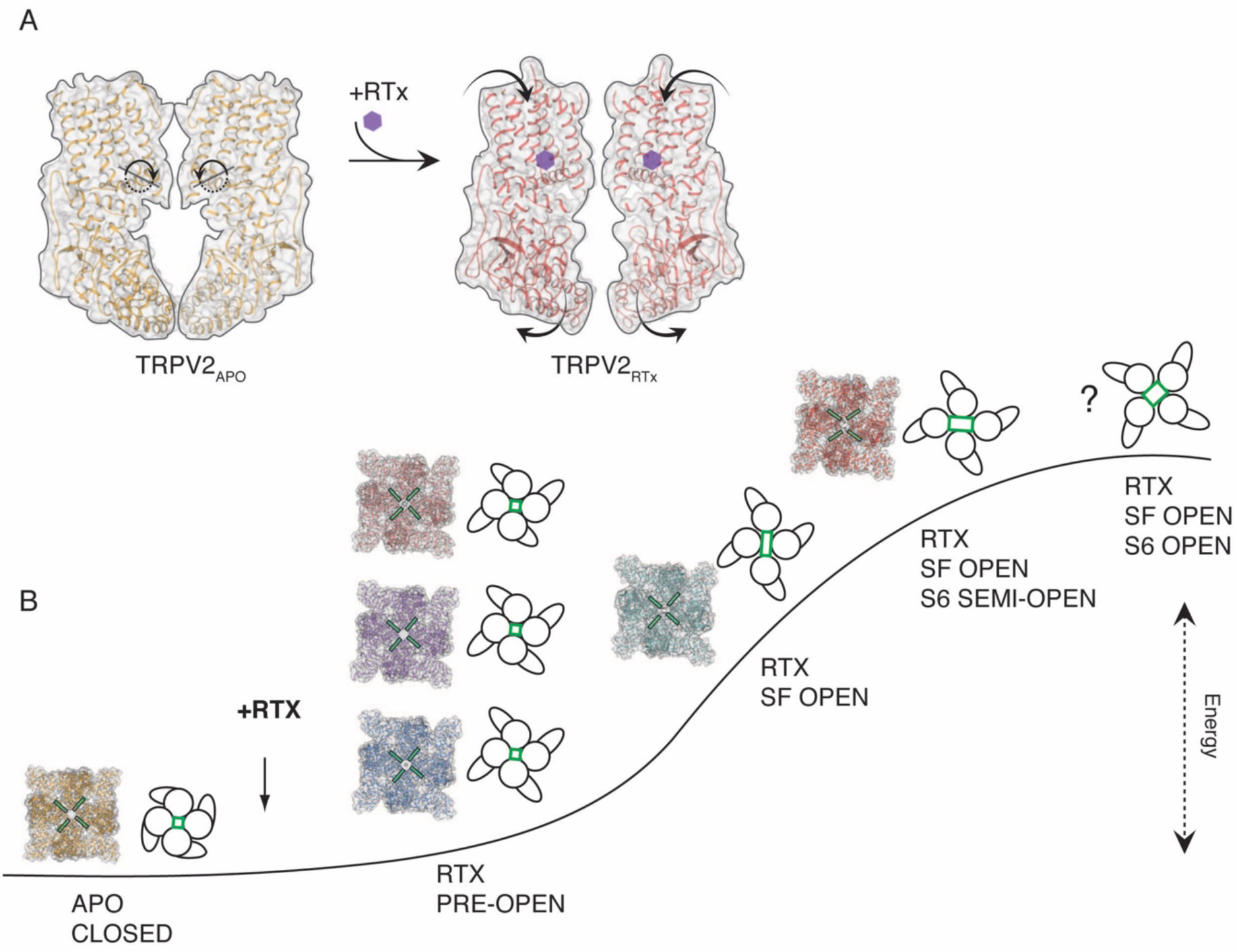
Conformational states associated with RTx-mediated gating of TRPV2. **A,** TRPV2 subunit rotation upon binding of RTx. Rotation axis and direction is indicated in dashed line and circular arrow in apo TRPV2 (left). The rotation results in contraction of the TM domains and widening of the cytoplasmic assembly (right). **B,** Hypothetical trajectory of TRPV2 gating with associated conformational states. Upon addition of RTx, TRPV2 first enters low-energy pre-open states that are characterized by rotation, widening and symmetry breaking in the ARD (TRPV2_RTx-APOL_ 1-3, models shown in cartoon and surface representation). In the next step, the channel assumes C2 symmetric state with an open SF gate, but closed common (S6) gate (TRPV2_RTx-XTAL,_ model shown in cartoon and surface representation). This is followed by a less C2 symmetric state with an open SF gate and semi-open S6 gate (TRPV2_RTx-ND_, model shown in cartoon and surface representation). Finally, we propose that the channel assumes a high-energy fully open state that is C4 symmetric but might have C2 symmetry in the SF gate. The SF gate is indicated in green in models and cartoons.

Two-fold symmetry is a well-stablished feature of mammalian Na^+^ selective Two Pore Channels (TPCs) and Voltage Gated Sodium channels (Na_V_)^39–42^. Interestingly, the arrangement of pore helices in TRPV2_RTx-ND_ resembles that observed in TPC and Na_V_ (Figure 6) and the selectivity filters in all three channels form a “coin-slot”^43^ opening. However, while the selectivity filters of TPC and Na_V_ remain static during channel gating in order to maintain the structure necessary for Na^+^ selectivity, the SF gate of TRPV2 displays a large degree of plasticity. Moreover, the two-fold symmetry observed in TRPV2 is unique in that it arises in response to conformational changes in the TM domains induced by ligand binding. By contrast, the two-fold symmetry in TPC and Na_V_ stems from the arrangement of their respective homologous tandem repeats.

**Figure 6.**
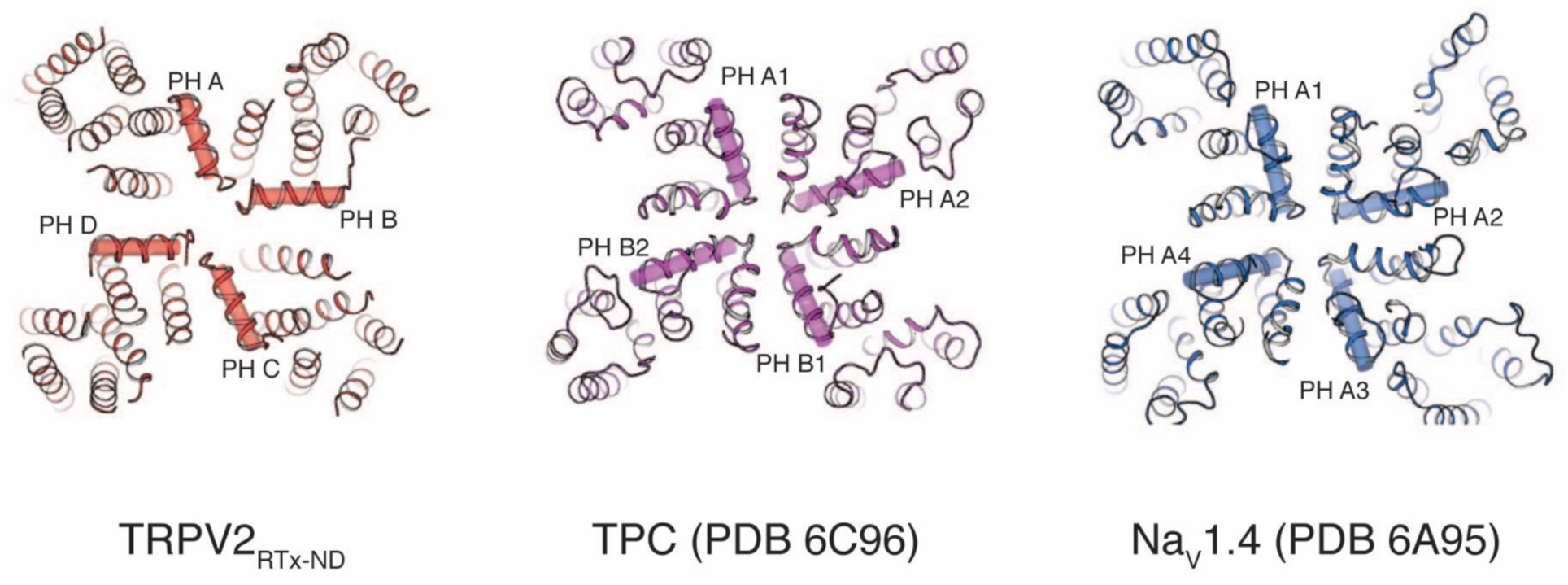
Comparison of TRPV2_RTx-ND_ (red), TPC (PDB 6C96, purple) and Na_V_1.4 (PDB 6A95, blue). Top view, pore helices are indicated.

## Methods

### Protein expression and purification

The construct for the RTx sensitive, full-length rabbit TRPV2 (TRPV2_RTx_) was prepared by introducing four point mutations (F470S, L505M, L508T and Q528E) into the synthesized full-length rabbit TRPV2 gene^23^. The construct was cloned into a pFastBac vector with a C-terminal FLAG affinity tag and used for baculovirus production according to manufacturers’ protocol (Invitrogen, Bac-to-Bac). The protein was expressed by infecting Sf9 cells with baculovirus at a density of 1.3M cells ml^-1^ and incubating at 27° C for 72 hours in an orbital shaker. Cell pellets were collected after 72 hours and resuspended in buffer A (50 mM TRIS pH8, 150 mM NaCl, 2 mM CaCl_2_, 1 μg ml^−1^ leupeptin, 1.5 μg ml^−1^ pepstatin, 0.84 μg ml^−1^ aprotinin, 0.3 mM PMSF, 14.3 mM β-mercapto ethanol, and DNAseI) and broken by sonication (3×30 pulses).

For the amphipol-reconstituted TRPV2 (TRPV2_RTx-APOL_) sample, the lysate was supplemented with 40 mM Dodecyl β-maltoside (DDM, Anatrace), 4 mM Cholesteryl Hemisuccinate (CHS, Anatrace) and 2 μM RTx and incubated at 4° C for 1 hour. Insoluble material was removed by centrifugation (8,000g, 30 minutes), and anti-FLAG resin was added to the supernatant for 1 hour at 4° C.

After binding, the anti-FLAG resin was loaded onto a Bio-Rad column and a wash was performed with 10 column volumes of Buffer B (50 mM TRIS pH8, 150 mM NaCl, 2 mM CaCl_2_, 1 mM DDM, 0.1 mM CHS, 0.1 mg ml^−1^ 1,2-dimyristoyl-*sn*-glycero-3-phosphocholine (DMPC, Avanti Polar Lipids), 2 μM RTx) before elution in 5 column volumes of buffer C (50 mM TRIS pH8, 150 mM NaCl, 2 mM CaCl_2_, 1mM DDM, 0.1 mM CHS, 0.1 mg ml^−1^ DMPC, 2 μM RTx, 0.1 mg ml^-1^ FLAG peptide).

The eluate was concentrated and further purified by gelfiltration on a Superose 6 column. The peak fractions were collected, mixed with Amphipol A8-35 (Anatrace) in a 1:10 ratio and incubated for 4 hours at 4° C. Subsequently, Bio-Beads SM-2 (Biorad) were added to a 50 mg ml^-1^ concentration and incubated at 4° C overnight to remove detergent.

After reconstitution, the protein was subjected to a second round of gelfiltration on a Superose 6 column in buffer D (50 mM TRIS pH8, 150 mM NaCl, 2 μM RTx), the peak fractions were collected and concentrated to 2-2.5 mg ml^-1^ for cryo-EM.

For the nanodisc reconstituted TRPV2 (TRPV2_RTx-ND_), the lysate was supplemented with 40 mM Dodecyl β-maltoside (DDM, Anatrace) and 2 μM RTx and incubated at 4° C for 1 hour. The solution was cleared by centrifugation (8,000g, 30 minutes), and anti-FLAG resin was added to the supernatant for 1 hour at 4° C.

After binding, the anti-FLAG resin was loaded onto a Bio-Rad column and a wash was performed with 10 column volumes of Buffer B_noCHS_ (50 mM TRIS pH8, 150 mM NaCl, 2 mM CaCl_2_, 1 mM DDM, 0.1 mg ml^−1^ DMPC, 2 μM RTx) before elution in 5 column volumes of buffer C_noCHS_ (50 mM TRIS pH8, 150 mM NaCl, 2 mM CaCl_2_, 1mM DDM, 0.1 mg ml^−1^ DMPC, 2 μM RTx, 0.1 mg ml^-1^ FLAG peptide).

The eluate from the anti-FLAG resin was concentrated to ~ 500 μl. A 10 mg ml^-1^ 3:1:1 mixture of lipids 1-palmitoyl-2-oleoyl-*sn*-glycero-3-phosphocholine (POPC), 1-palmitoyl-2-oleoyl-*sn*-glycero-3-phosphoethanolamine (POPE), 1-palmitoyl-2-oleoyl-*sn*-glycero-3-phospho-(1’-*rac*-glycerol) (POPG) was dried under argon, resuspended in 1 ml 50 mM Tris pH8, 150 mM NaCl and clarified by extrusion, before being incubated for 1 hour with 10mM DDM. The membrane scaffold protein MSP2N2 was prepared as previously described^44^. The concentrated TRPV2 was combined with MSP2N2 and the prepared lipid mixture in a 1:3:200 ratio and incubated at 4° C for 1 hour. After the initial incubation, 50 mg ml^-1^ Bio-Beads SM-2 were added and the mixture was incubated for another hour at 4° C, following which the reconstitution mixture was transferred to a fresh batch of Bio-Beads SM-2 at 50 mg ml^-1^ and incubated overnight at 4° C. Finally, the reconstituted channels were subjected to gelfiltration on Superose 6 in buffer D, the peak fractions collected and concentrated to 2-2.5 mg ml^-1^ for cryo-EM.

### Cryo-EM sample preparation

TRPV2_RTx-APOL_ and TRPV2_RTx-ND_ were frozen using the same protocol. Before freezing, the concentrated protein sample was supplemented with 300 μM RTx and incubated ~30 minutes at 4° C. 3 μl sample was dispensed on a freshly glow discharged (30 seconds) UltrAuFoil R1.2/1.3 300-mesh grid (Electron Microscopy Services), blotted for 3 seconds with Whatman No. 1 filter paper using the Leica EM GP2 Automatic Plunge Freezer at 23° C and > 85% humidity and plunge-frozen in liquid ethane cooled by liquid nitrogen.

### Cryo-EM data collection

Data for both TRPV2_RTx-APOL_ and TRPV2_RTx-ND_ was collected using the Titan Krios transmission electron microscope (TEM) operating at 300 keV using a Falcon III Direct Electron Detector operating in counting mode at a nominal magnification of 75,000x corresponding to a physical pixel size of 1.08 Å/pixel.

For the TRPV2_RTx-APOL_ 1293 movies (30 frames/movie) were collected using a 60 second exposure with an exposure rate of ~0.8 e^-^/pixel/s, resulting in a total exposure of 42 e^-^/Å^2^ and a nominal defocus range from −1.25 µm to −3.0 µm.

For TRPV2_RTx-ND_, 2254 movies were collected (30 frames/movie) with 60 second exposure and exposure rate of ~0.8 e^-^/pixel/s. The total exposure was of 42 e^-^/Å^2^ and a nominal defocus range from −1.25 µm to −3.0 µm.

### Reconstruction and refinement

*TRPV2_RTx-APOL_* MotionCor2^45^ was used to perform motion correction and dose-weighting on 1293 movies. Unweighted summed images were used for CTF determination using GCTF^46^. Following motion correction and dose-weighting and CTF determination, micrographs which contained Figure of Merit (FoM) values of < 0.12 and astigmatism values > 400 were removed, leaving 1207 micrographs for further analysis. An initial set of 1660 particles was picked manually and subjected to reference-free 2D classification (k= 12, T=2) which was used as a template for automatic particle picking from the entire dataset (1207 micrographs). This yielded a stack of 580,746 particles that were binned 4 × 4 (4.64 Å/pixel, 64 pixel box size) and subjected to reference-free 2-D classification (k=58, T=2) in RELION 3.0^47^. Classes displaying the most well-defined secondary structure features were selected (470,760 particles) and an initial model was generated from the 2D particles using the Stochastic Gradient Descent (SGD) algorithm as implemented in RELION 3.0. 3D auto-refinement in RELION 3.0 was performed on the 470,760 particles with no symmetry imposed (C1), using the initial model, low-pass filtered to 30 Å, as a reference map. This resulted in an 8.9 Å 3D reconstruction, which was then used for re-extraction and re-centering of 2 × 2 binned particles (2.16 Å/pixel, 128 pixel box size). 3D classification (k=4, T=8) without imposed symmetry (C1) was performed on the extracted particles, using a soft mask calculated from the full molecule. Classes 2-4 (90,862, 109,623 and 101,570 particles, respectively) all possessed well-defined secondary structure, but visual inspection of the maps suggested that the classes represented distinct conformational states. Therefore, each class was processed separately. For each class, the particles were extracted and unbinned (1.08 Å/pixel, 256 pixel box size), and soft masks calculated. 3D auto-refinement of the individual classes without symmetry imposed (C1) yielded 4.7 Å (class 2), 3.6 Å (class 3) and 3.2 Å (class 4) 3D reconstructions. Inspection of these volumes revealed that classes 2 and 3 adopted two-fold (C2) symmetry, while class 4 was four-fold symmetric (C4). Particles from class 2 were subjected to particle movement and dose-weighting using the “particle polishing” function as implemented in RELION 3.0. The shiny particles were input into 3D auto-refinement with a soft mask and C2 symmetry applied, resulting in a 4.19 Å reconstruction (TRPV2_RTx-APOL_ 3). Similarly, particles from class 3 were subjected to polishing, and the following 3D auto-refinement with a soft mask and C2 symmetry applied resulted in a 3.3 Å final reconstruction (TRPV2_RTx-APOL2_). Particles from class 4 were first subjected to CTF refinement using the “CTF refine” feature in RELION 3.0. Particle polishing was then performed, followed by 3D auto-refinement with a soft mask and C4 symmetry applied, yielding a 2.91 Å reconstruction (TRPV2_RTx-APOL 1_). All resolution estimates were based on the gold-standard FSC 0.143 criterion^48,49^.

*TRPV2_RTx-ND_* The 2254 collected movies were subjected to motion correction and dose-weighting (MotionCor2) and CTF estimation (GCTF) in RELION 3.0. Micrographs with FoM values < 0.13 and astigmatism values > 400 were removed, resulting in a selection of 1580 good micrographs. From these, 2015 particles were picked manually, extracted (1×1 binned, 1.08 Å/pixel, 256 pixel box size) and subjected to reference-free 2D classification (k=12, T=2) that was used as a template for autopicking. This resulted in a 1,407,292 stack of particles that were binned 4×4 (4.32 Å/pixel, 64 pixel box size) and subjected to reference-free 2D classification (k=100, T=2). Classes exhibiting the most well-defined secondary structure features were selected, resulting in 482,602 particles. These were re-extracted (2×2 binned, 2.16 Å/pixel, 128 pixel box size) and put into 3D auto-refinement, using the previously obtained map of apo TRPV2 (EMD-6455) filtered to 30 Å as a reference with no symmetry applied (C1). The 3D auto-refinement yielded a 5.4 Å reconstruction. The particles were then subjected to 3D classification (k=6, T=8), with a soft mask and the 5.4 Å volume as a reference without imposed symmetry (C1). Only two of the six classes (classes 1 and 6) contained significant density in the TM domains. They were selected (112,622 particles), re-extracted, re-centered and unbinned (1.08 Å/pixel, 256 pixel box size) before being input into 3D auto-refinement without symmetry imposed (C1) and with a soft mask and the previous 5.4 Å reconstruction filtered to 30 Å as a reference. The 3D auto-refinement resulted in a 4.12 Å map, which was then subjected to Bayesian particle polishing. 3D auto-refinement was then performed on the resulting shiny particles with no symmetry applied (C1), resulting in a 4 Å reconstruction. The particles were then subjected to CTF refinement, yielding a 3D reconstruction resolved to 4 Å (C1). However, visual inspection of the map revealed a strong tendency towards two-fold symmetry. Therefore, 3D auto-refinement was repeated with C2 symmetry applied, resulting in a map resolved to 3.84 Å as estimated by gold-standard FSC 0.143 criterion

### Model building

The TRPV2_RTx-APOL_ and TRPV2_RTx-ND_ models were built into the cryo-EM electron density map in Coot^50^, using the structures of TRPV2 (PDB 5AN8 and 6BWM) as templates. The structures were real-space refined in Coot, and iteratively refined using the phenix.real_space_refine as implemented in the Phenix suite^51^. Structures were refined using global minimization and rigid body, with high weight on ideal geometry and secondary structure restraints. The Molprobity server^52^ (http://molprobity.biochem.duke.edu/) was used to identify problematic areas, which were subsequently manually rebuilt. The radius of the permeation pathways was calculated using HOLE^53^. All analysis and structure illustrations were performed using Pymol (The PyMOL Molecular Graphics System, Version 2.0) and UCSF Chimera^54^.

## Acknowledgements

Cryo-EM data were collected at the Shared Materials Instrumentation Facility at Duke University as part of the Molecular Microscopy Consortium, and screening was performed at NIEHS. We thank Alberto Bartesaghi for a pre-processing interface.

## Funding

This work was supported by the National Institutes of Health (R35NS097241 to S.-Y.L.) and by the National Institutes of Health Intramural Research Program; US National Institute of Environmental Health Science (ZIC ES103326 to M.J.B). The EM maps and atomic models have been deposited with the Electron Microscopy Data Bank (accession numbers ###, ###, ###, and ###) and the Protein Data Back (entry codes ###, ###, ###, and ###), respectively.

## Competing Interests

The authors declare no competing interests.

---

**Figure Supplement 1.**
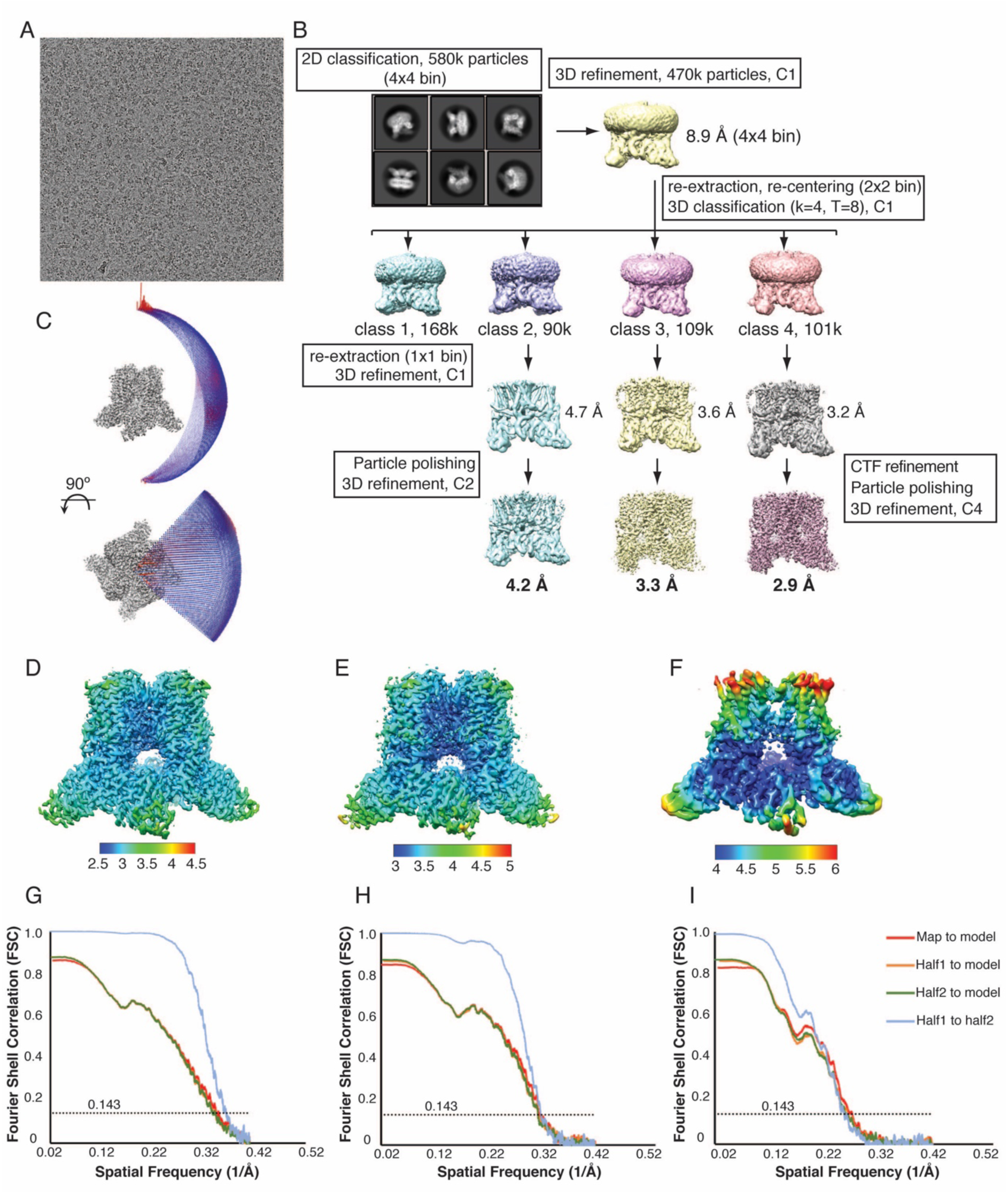
Cryo-EM data collection and processing, TRPV2_RTx-APOL._ **A**, Representative micrograph from the TRPV2_RTx-APOL_ dataset. **B**, 3D reconstruction workflow resulting in 3 distinct TRPV2_RTx-APOL_ structures. **C**, Euler plot distribution. Red regions signify the best represented views. **D-F**, Local resolution estimates calculated in Relion for TRPV2_RTx-APOL_ 1 **(D)**, TRPV2_RTx-APOL_ 2 **(E)**, TRPV2_RTx-APOL_ 3 **(F)**. **G-I**, FSC curves calculated between the half maps (blue), atomic model and the final map (red), and between the model and each half-map (orange and green) for TRPV2_RTx-APOL_ 1 **(G)**, TRPV2_RTx-APOL_ 2 **(H)**, TRPV2_RTx-APOL_ 3 **(I)**.

**Figure Supplement 2.**
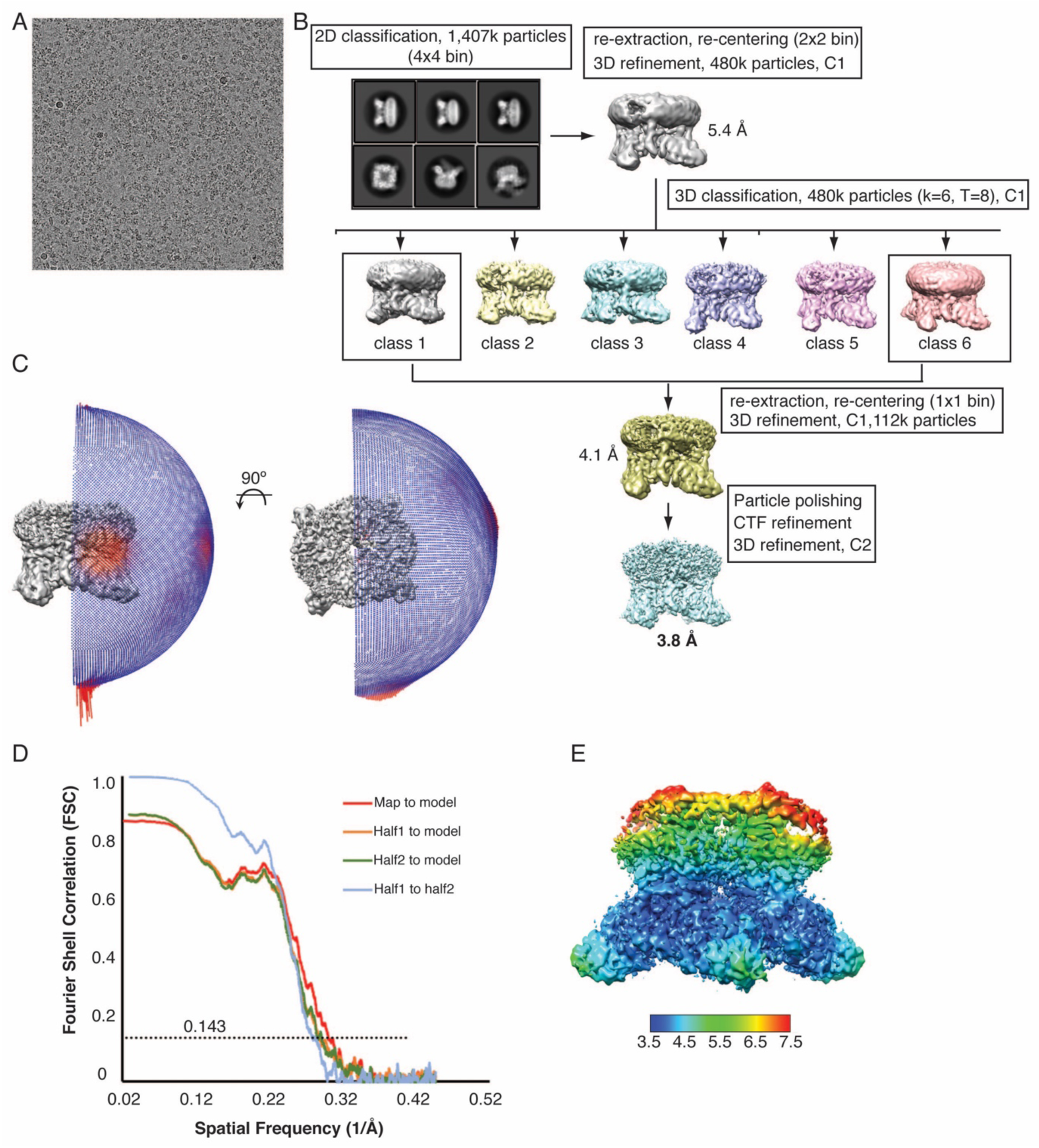
Cryo-EM data collection and processing, TRPV2_RTx-ND_. **A,** Representative micrograph from the collected TRPV2_RTx-ND_ dataset. **B,** 3D reconstruction workflow. **C,** Euler distribution plot. Red regions indicate best represented views. **D,** FSC curves calculated between the half maps (blue), atomic model and the final map (red), and between the model and each half-map (orange and green). **E,** Local resolution estimate, calculated in Relion.

**Figure Supplement 3.**
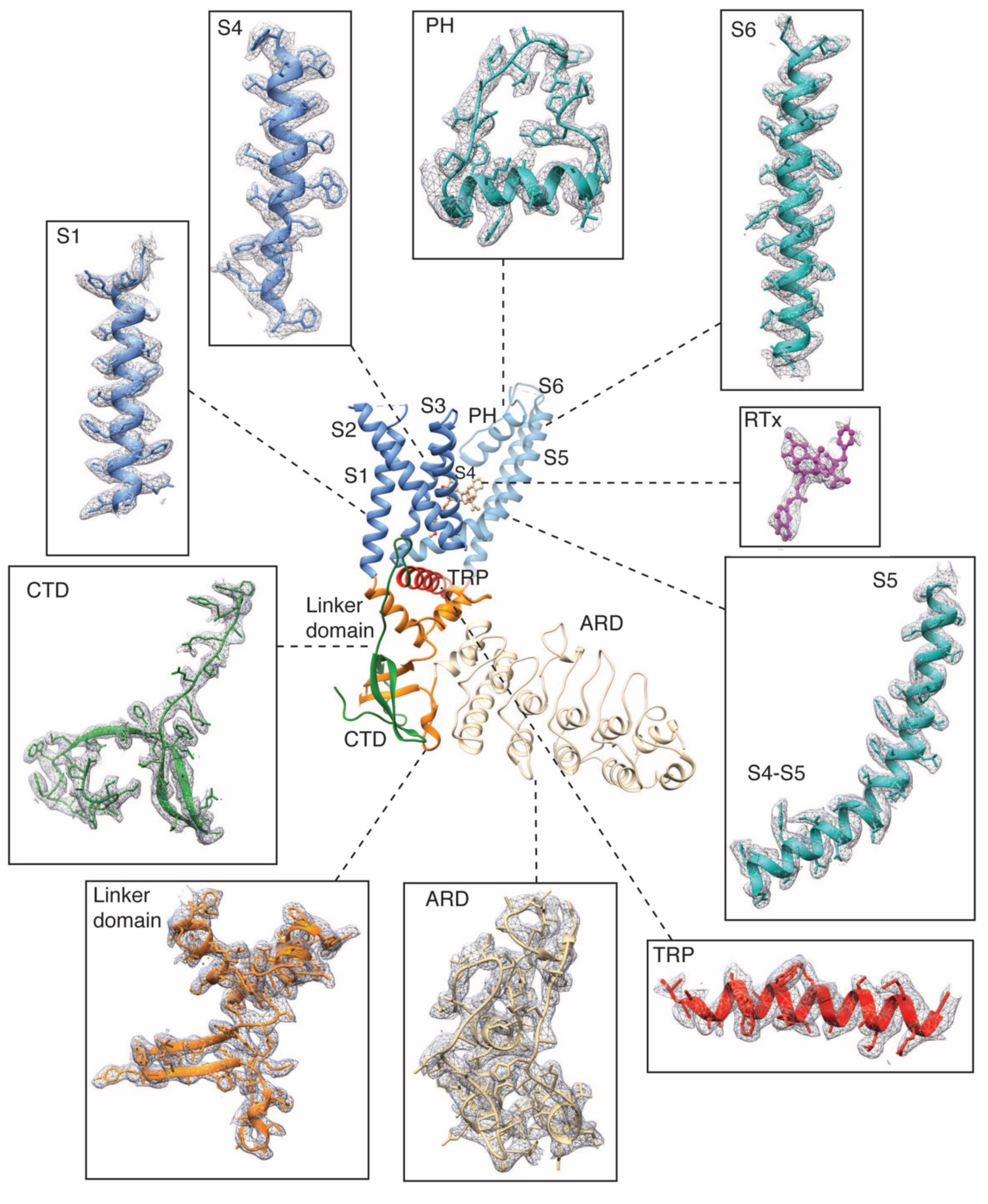
Representative electron densities in the TRPV2_RTx-APOL_ 1 cryo-EM map. Densities are contoured at level 0.06 and radius 2.

**Figure Supplement 4.**
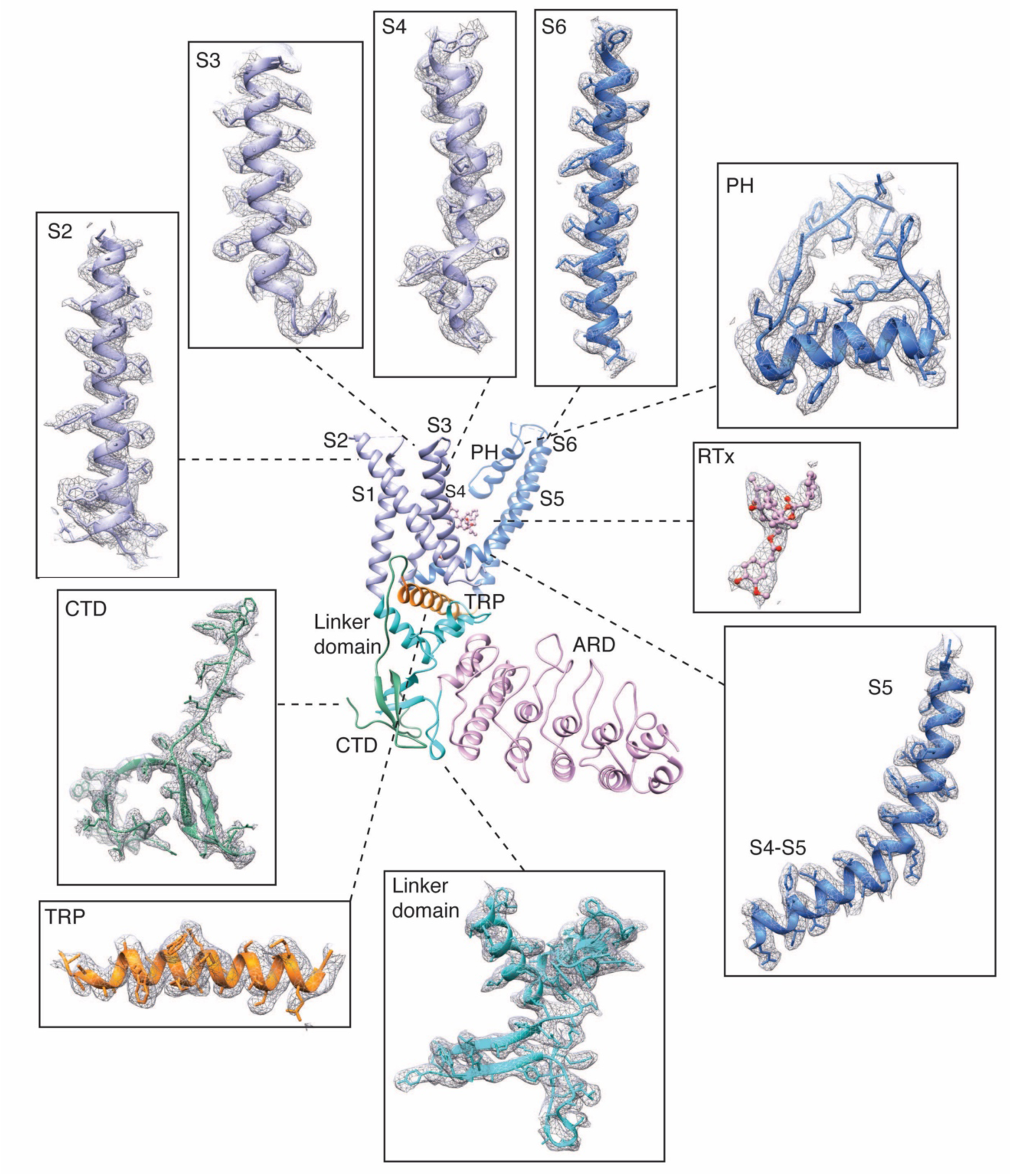
Representative electron densities in the TRPV2_RTx-APOL_ 2 cryo-EM map. Densities are contoured at level 0.06 and radius 2.

**Figure Supplement 5.**
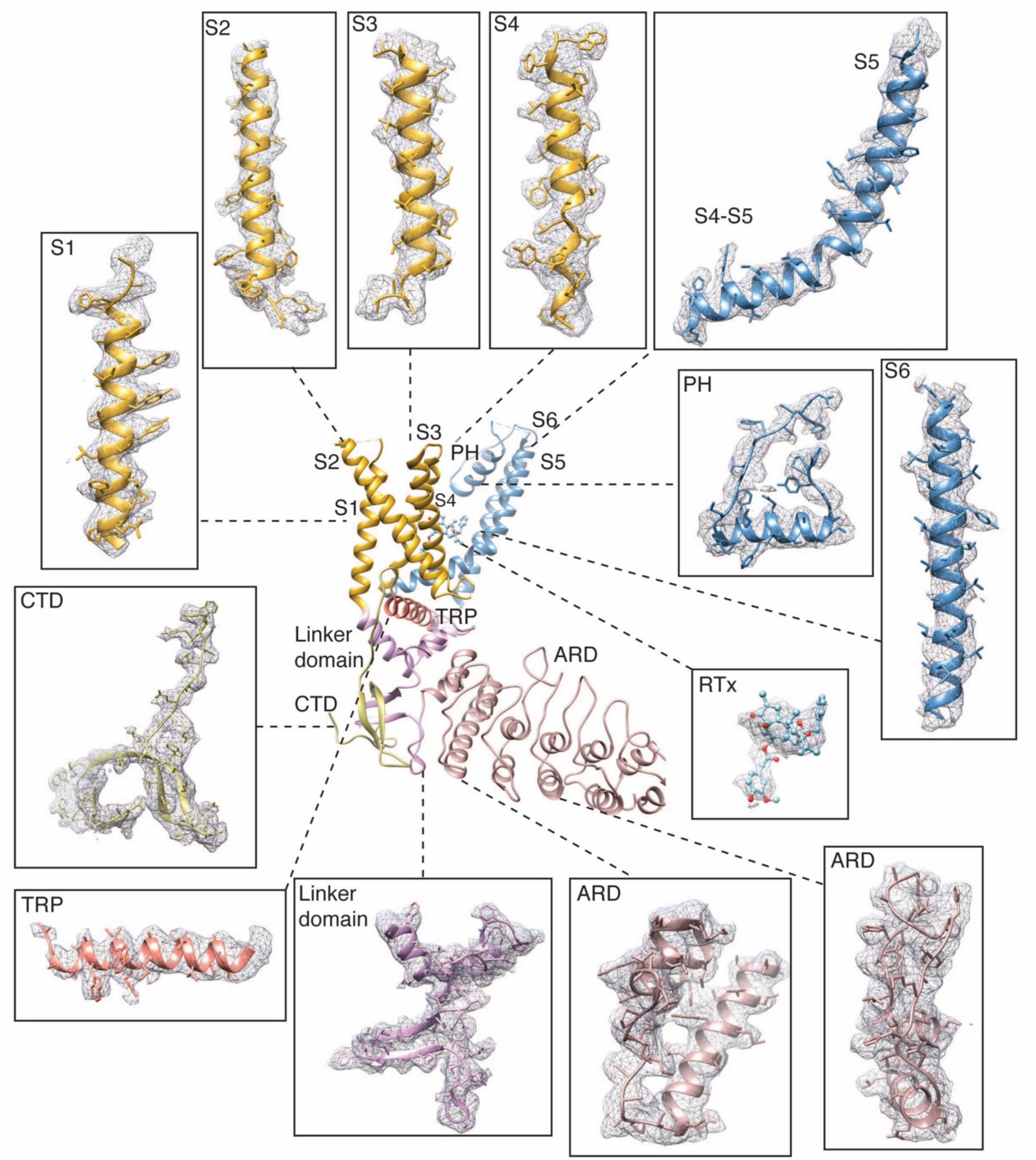
Representative electron densities in the TRPV2_RTx-APOL_ 3 cryo-EM map. Densities are contoured at level 0.02 and radius 2.

**Figure Supplement 6.**
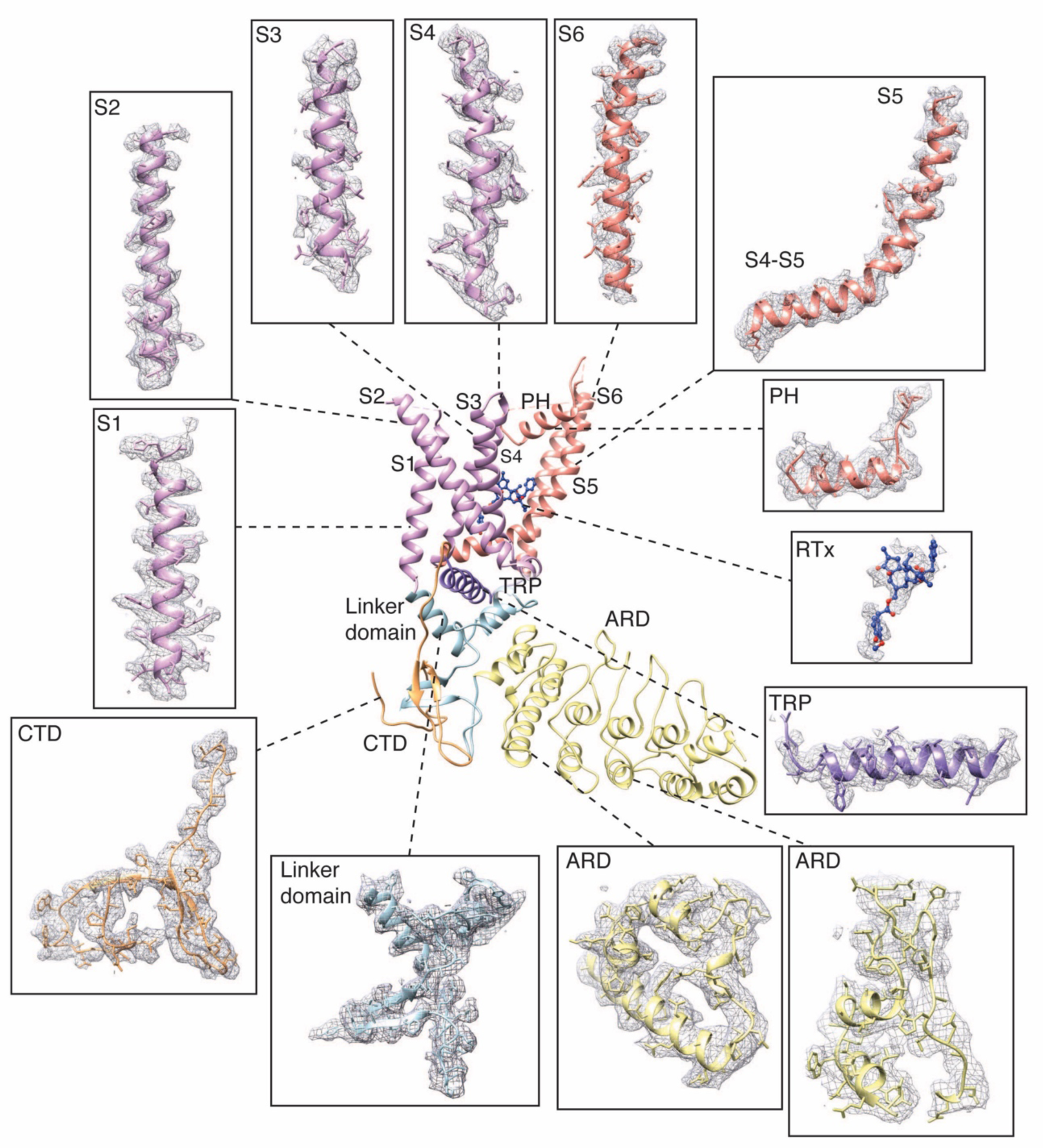
Representative electron densities in the TRPV2_RTx-ND_ cryo-EM map. Densities are contoured at level 0.015-0.03 and radius 2.

**Figure Supplement 7.**
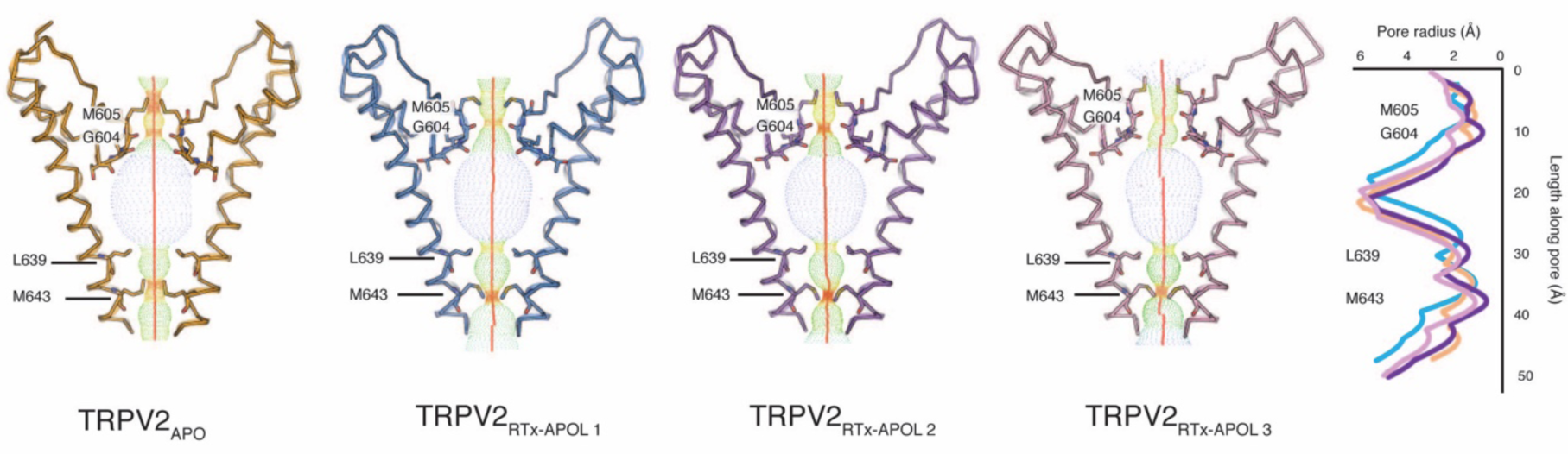
Pore comparison of TRPV2_APO_ (orange) and TRPV2_RTx-APOL_ 1-3 (blue, purple and salmon, respectively). HOLE profiles (dots and graph) indicate that both the selectivity filter and the common gates are closed in TRPV2_RTx-APOL_ 1-3.

**Figure Supplement 8.**
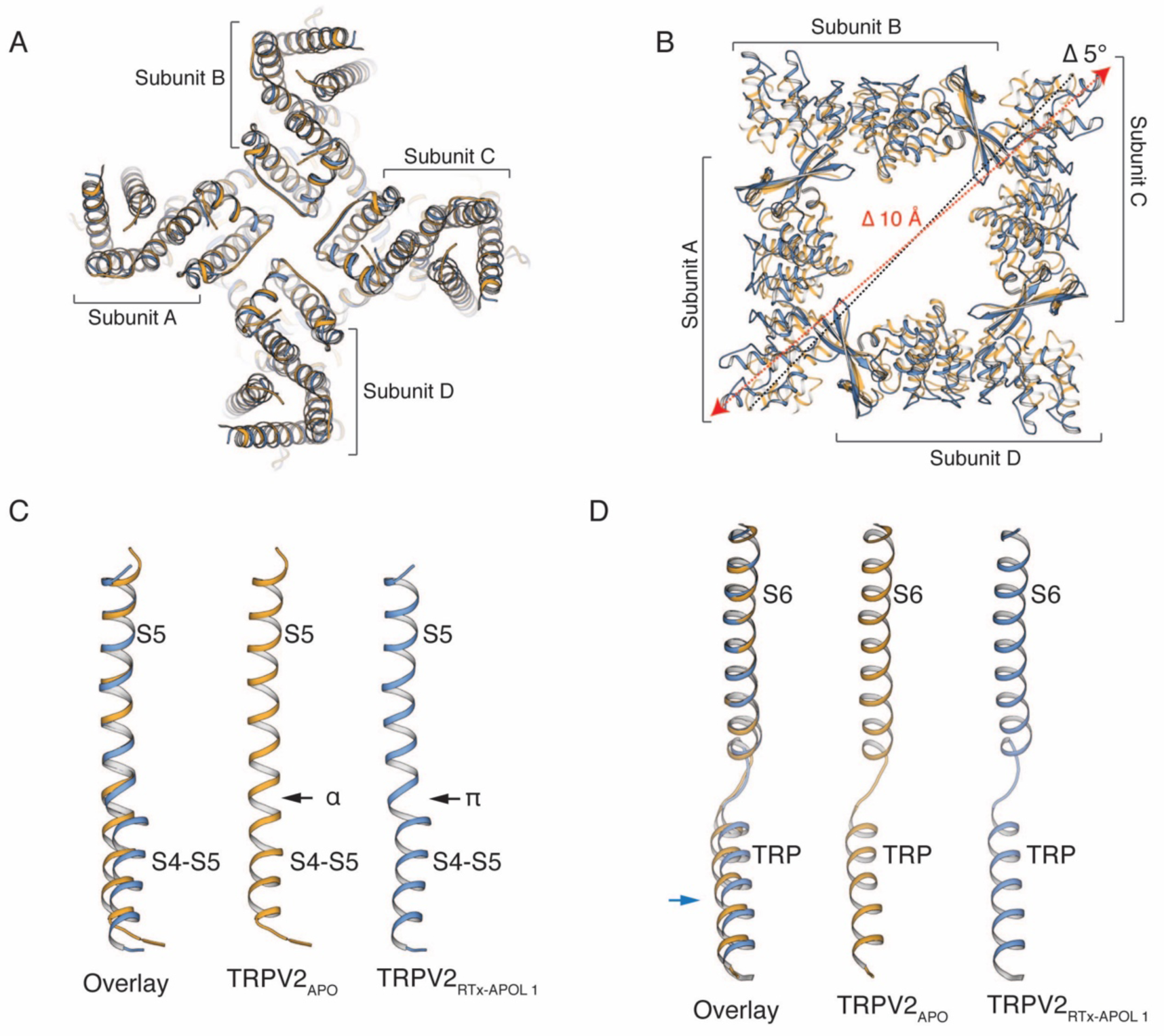
Comparison of TRPV2_RTx-APOL_ 1 (blue) and TRPV2_APO_ (orange). **A,** Overlay of the TM helices. Individual subunits are indicated. **B,** Top view of the cytoplasmic domains. The TMs are removed for ease of viewing. Distance measured between residues T100 in TRPV2_APO_ (black dotted line) and TRPV2_RTx-APOL_ 1 (red dotted line). The cytoplasmic assembly rotates by 5° and widens by 10Å in the presence of RTx. **C,** Overlay of S5 helices. In the presence of RTx, a π-helix is formed at the junction of the S4-S5 linker and the S5 helix changing the position of the S4-S5 linker. **D,** Overlay of S6 helices and the TRP domain. The TRP domain is displaced in the presence of RTx.

**Figure Supplement 9.**
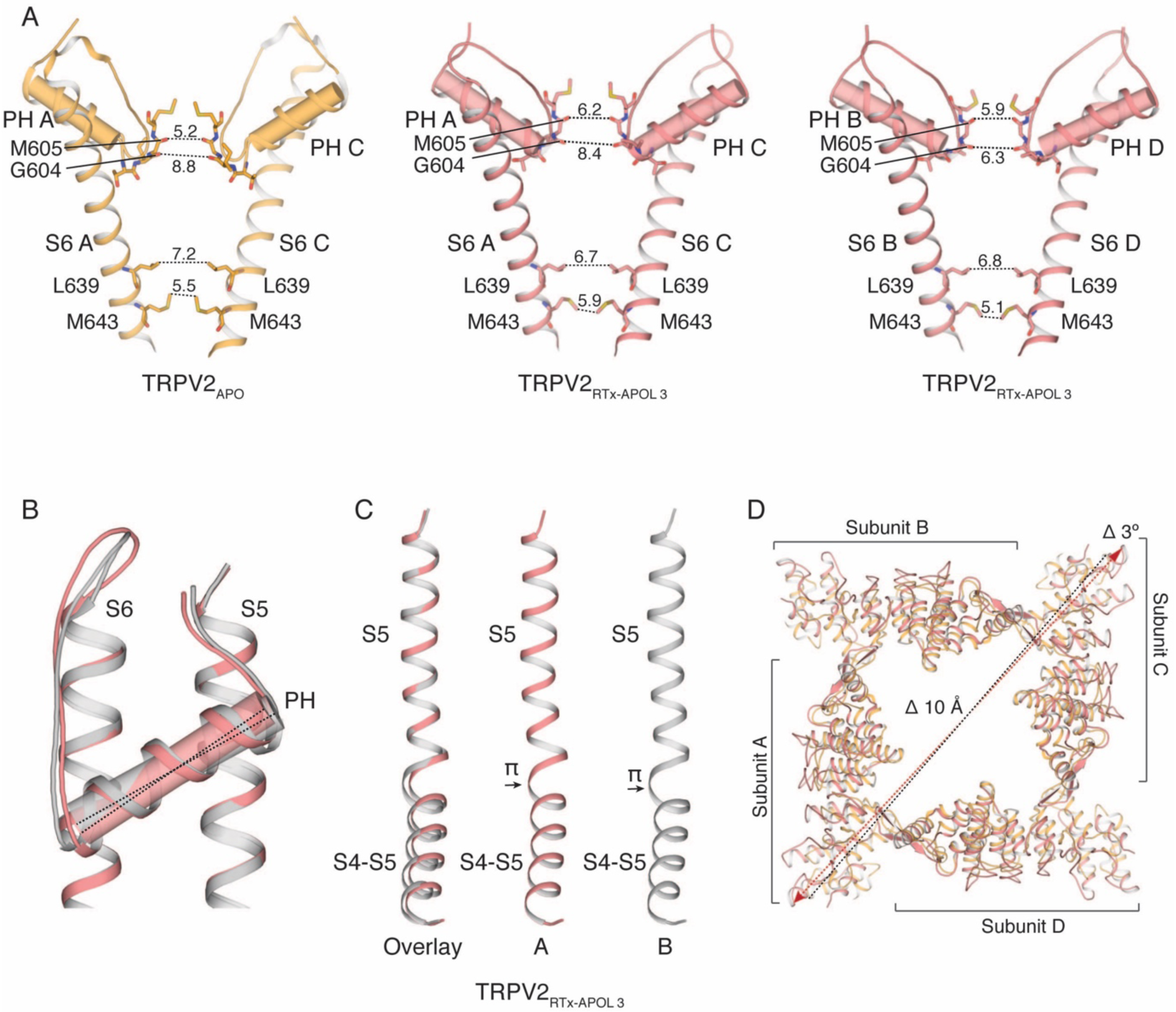
Two-fold symmetry in TRPV2_RTx-APOL_ 3 (salmon). **A,** Pore of the four-fold symmetric TRPV2_APO_ (orange) compared to the pore of the two-fold symmetric TRPV2_RTx-APOL_ 3 (salmon) subunits A and C (middle) and subunits B and D (right). **B,** Position of the pore helix in TRPV2_RTx-APOL_ 3 subunit A (salmon) compared to the subunit B (grey). **C,** Conformation of the S4-S5 linker in TRPV2_RTx-APOL_ 3 subunit A (salmon) compared to subunit B (grey). **D,** Comparison of TRPV2_RTx-APOL_ 3 (salmon) and TRPV2_APO_ (orange) ARD. The TMs are removed for ease of viewing. The dashed lines represent the distance between diagonally opposite residues T100 in TRPV2_APO_ (black line) and TRPV2_RTx-APOL_ 3 (red line). The ARD are rotated and expanded in TRPV2_RTx-APOL 3_.

**Figure Supplement 10.**
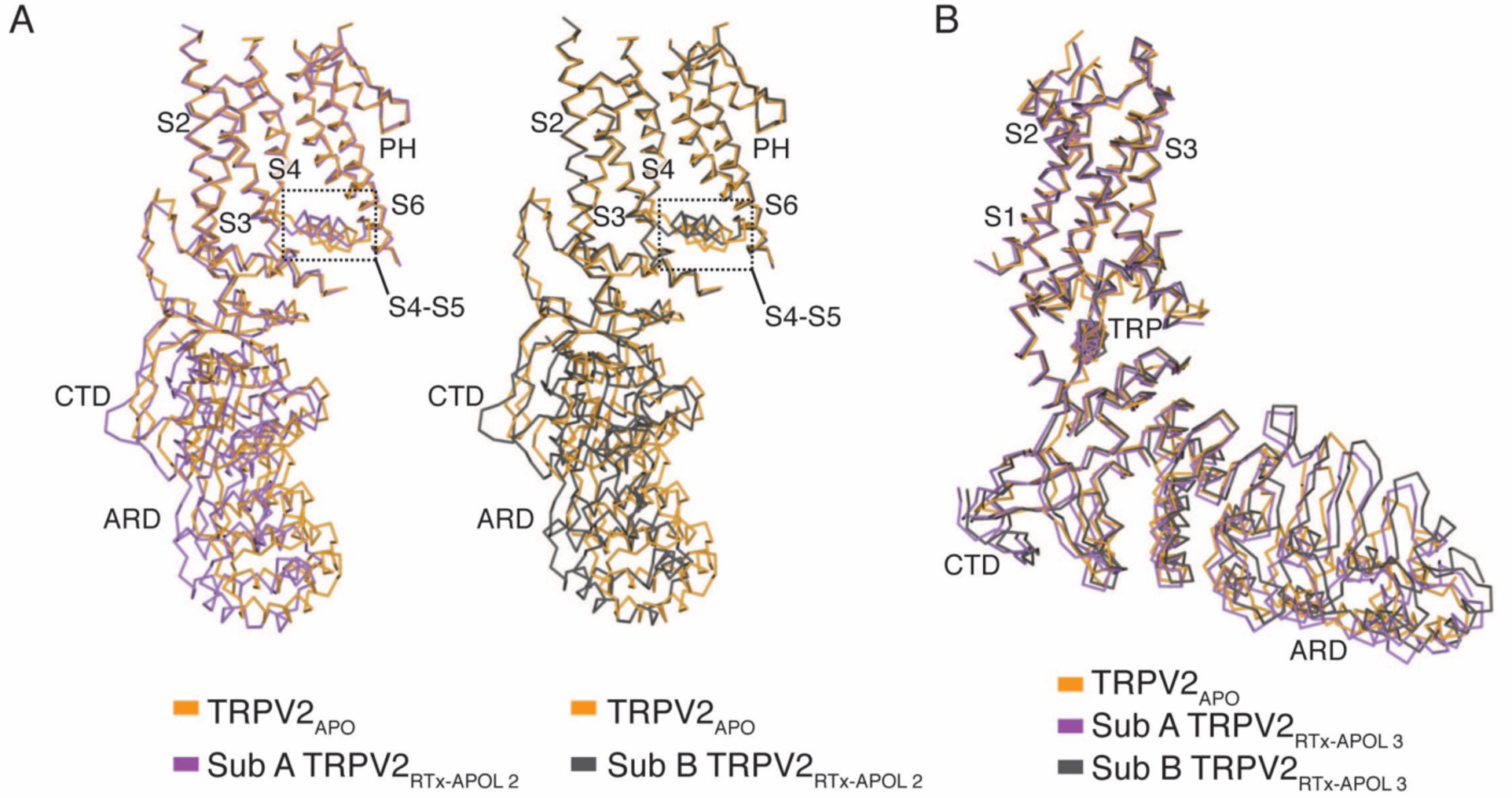
Symmetry breaking in the TRPV2_RTx-APOL_ 2-3 ARD. **A,** Two-fold symmetry in the ARD and S4-S5 linker of the TRPV2_RTx-APOL_ 2 structure. Subunit A (purple) overlaid with TRPV2_APO_ (orange) (left). Subunit B (purple)overlaid with TRPV2_APO_ (orange). In both subunits, TM domains are aligned but ARD and the S4-S5 linker (dashed line box) diverge. **B,** TRPV2_RTx-APOL_ 2 subunits A and B (purple) assume distinct conformations in the ARD.

**Figure Supplement 11.**
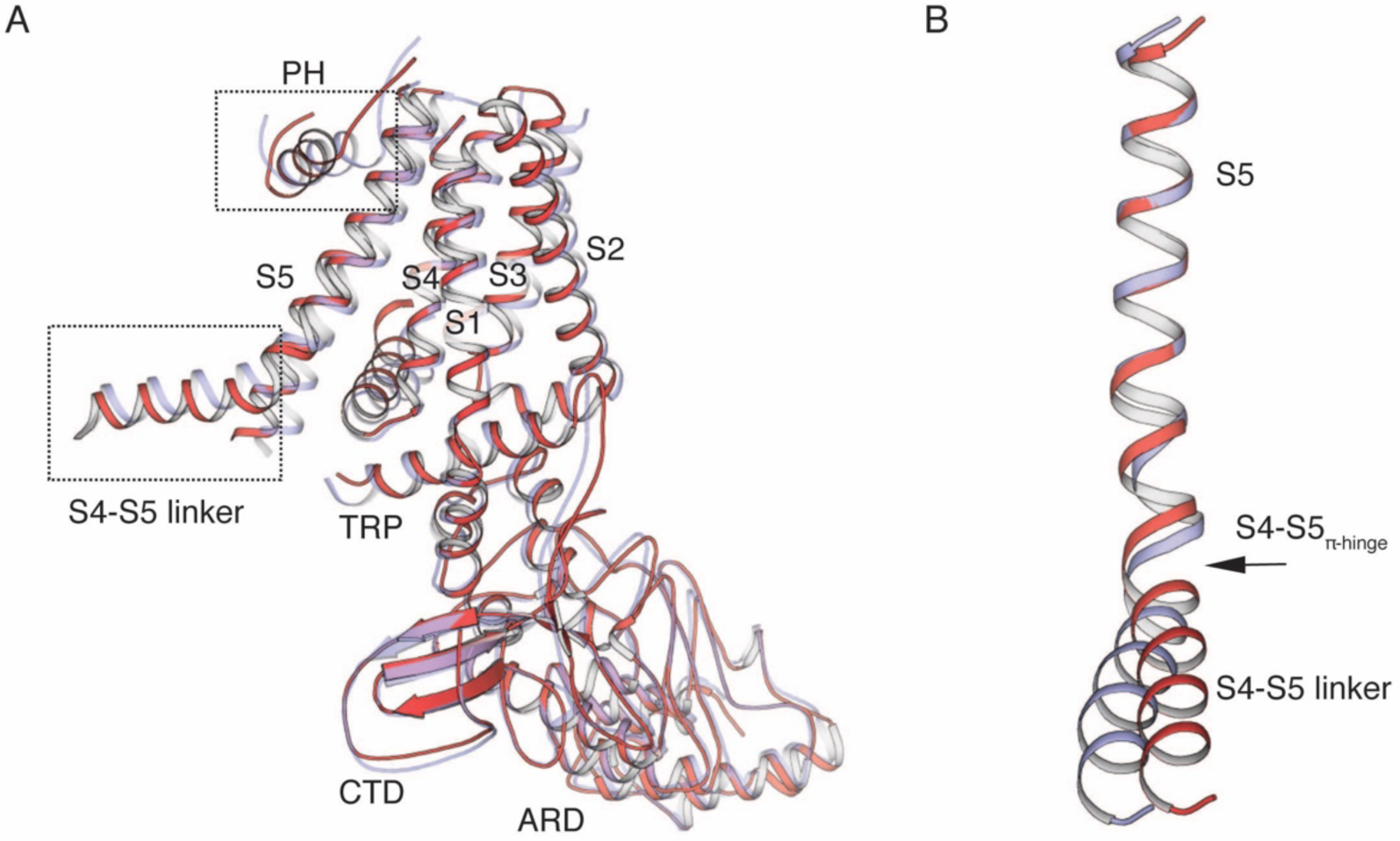
Comparison of TRPV2_RTx-ND_ subunits A (red) and B (violet). **A,** Overlay of the subunits. The regions that diverge from the overlay, the S4-S5 linker and the pore helix (PH), are indicated by a dashed line box. **B,** Overlay of S5 helices. The alignment diverges at the S4-S5 linker π-helix (S4-S5_π-hinge_) giving rise to different conformations of the S4-S5 linker in the two subunits.

**Figure Supplement 12.**
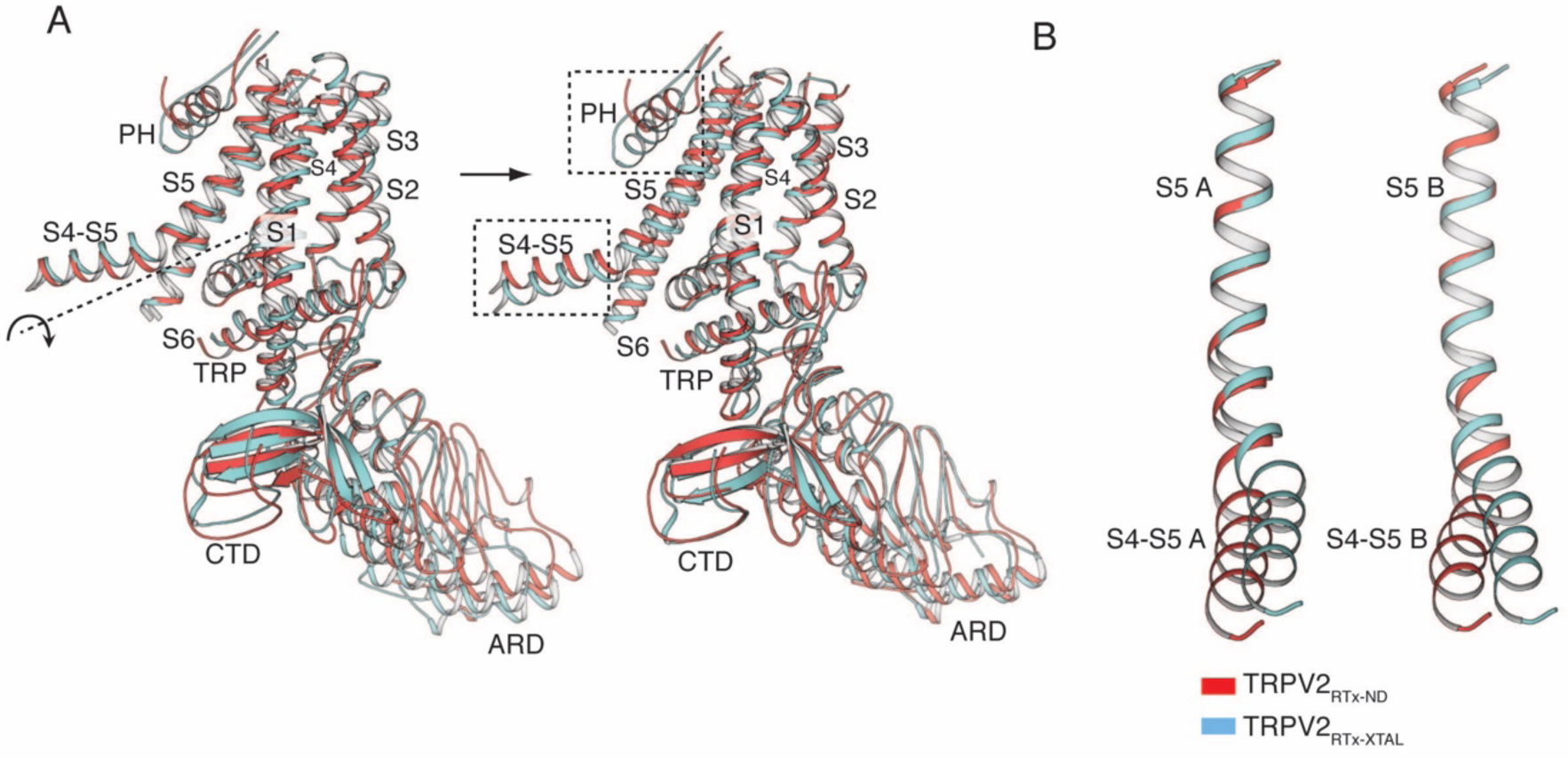
Comparison of subunits B in TRPV2_RTx-APOL_ (red) and TRPV2_RTx-XTAL_ (cyan). **A,** Rotation of subunit B from TRPV2_RTx-APOL_ around the S4-S5_π-hinge_ aligns it to subunit B from TRPV2_RTx-XTAL_. The S4-S5 linker and the PH (dashed box) diverge from the alignment. Rotation axis indicated with dashed line and arrow. **B,** Overlay of TRPV2_RTx_-APOL and TRPV2_RTx-XTAL_ S5 helices from subunit A (left) and subunit B (right) show that the S4-S5 linkers assume different conformations.

**Figure Supplement 13.**
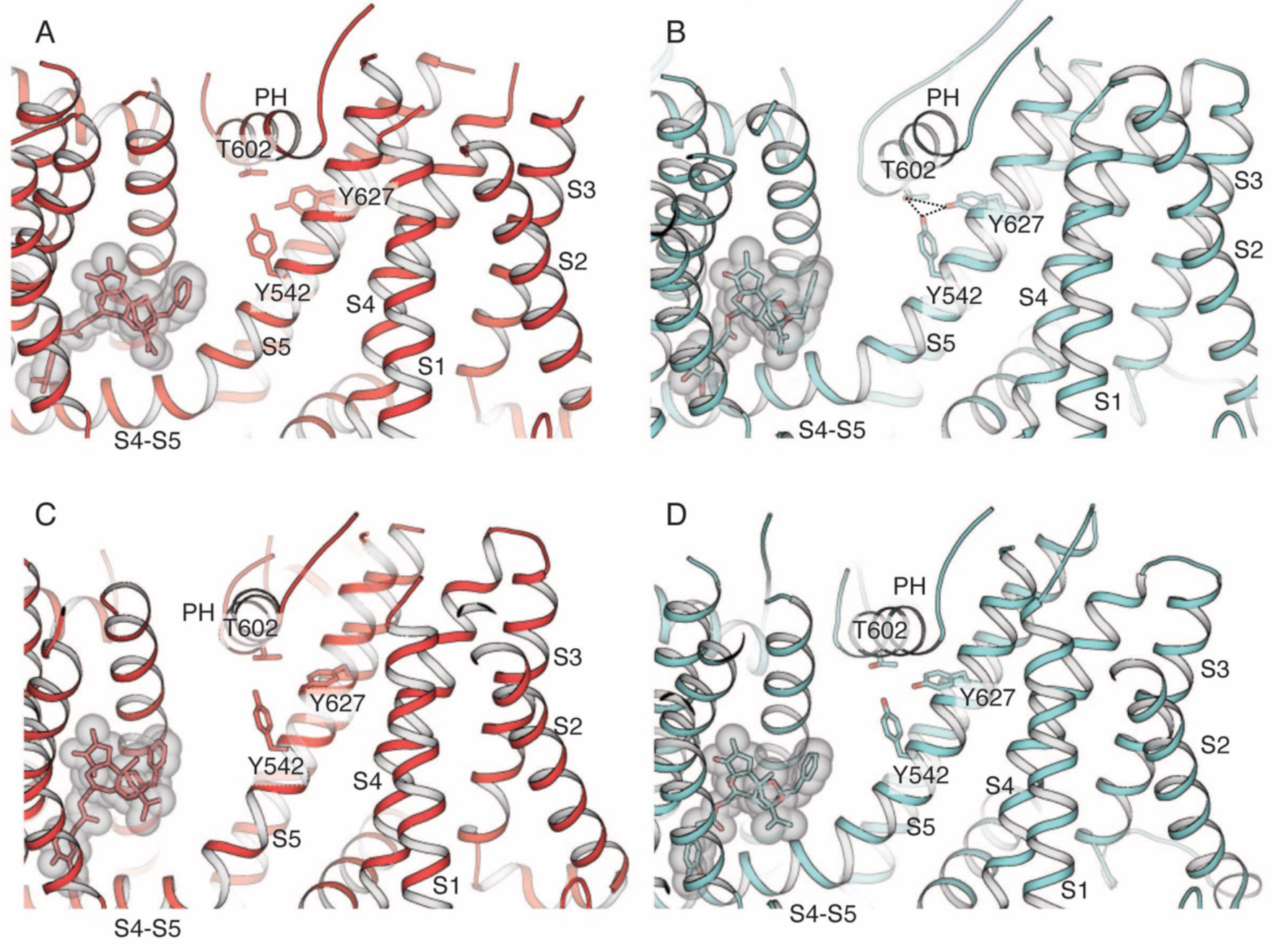
Interactions between the pore helix (PH) and S5 and S6. RTx is shown in stick and transparent surface representation. **A-B,** Side view of subunits A in TRPV2_RTx-ND_ **(A)** and TRPV2_RTx-XTAL_ **(B)**. The hydrogen bond triad (Y542-T602-Y627) is present in subunit A of TRPV2_RTx-XTAL_. The triad is broken in TRPV2_RTx-ND_. **C-D,** Side view of subunits B in TRPV2_RTx-ND_ **(C)** and TRPV2_RTx-XTAL_ **(D)**. The hydrogen bond triad is absent in both structures.

**Figure Supplement 14.**
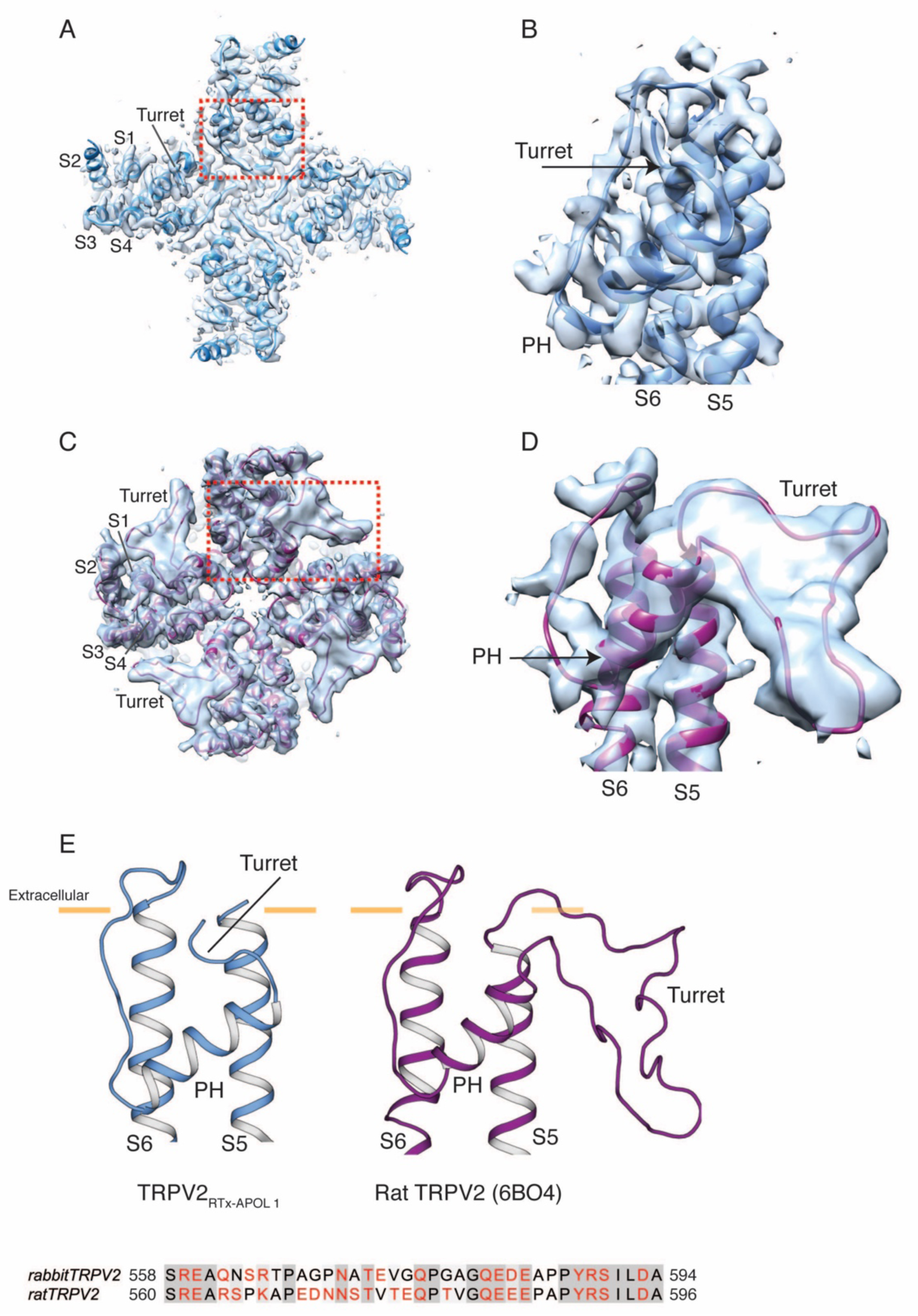
The pore turret in TRPV2_RTx-APOL_ 1 (blue) and rat TRPV2 (PDB 6BO4, purple). **A,** Top view of the map and model of TRPV2_RTx-APOL_ 1 with the pore domain indicated by dashed red box. **B,** Side view of the map and model of the pore domain in TRPV2_RTx-APOL_ 1. S5, S6, PH and pore turret are indicated. **C,** Top view of the map and model of rat TRPV2 with the pore domain indicated by dashed red box. **D,** Side view of the map and model of the pore domain in rat TRPV2. S5, S6, PH and pore turret are indicated. **E,** Position of the turret relative to the membrane (yellow lines) in TRPV2_RTx-APOL 1_ and rat TRPV2. The sequence of the turret shows conservation (gray boxes) and amino acids colored in red indicate charged or polar residues

**Figure Supplement 15.**
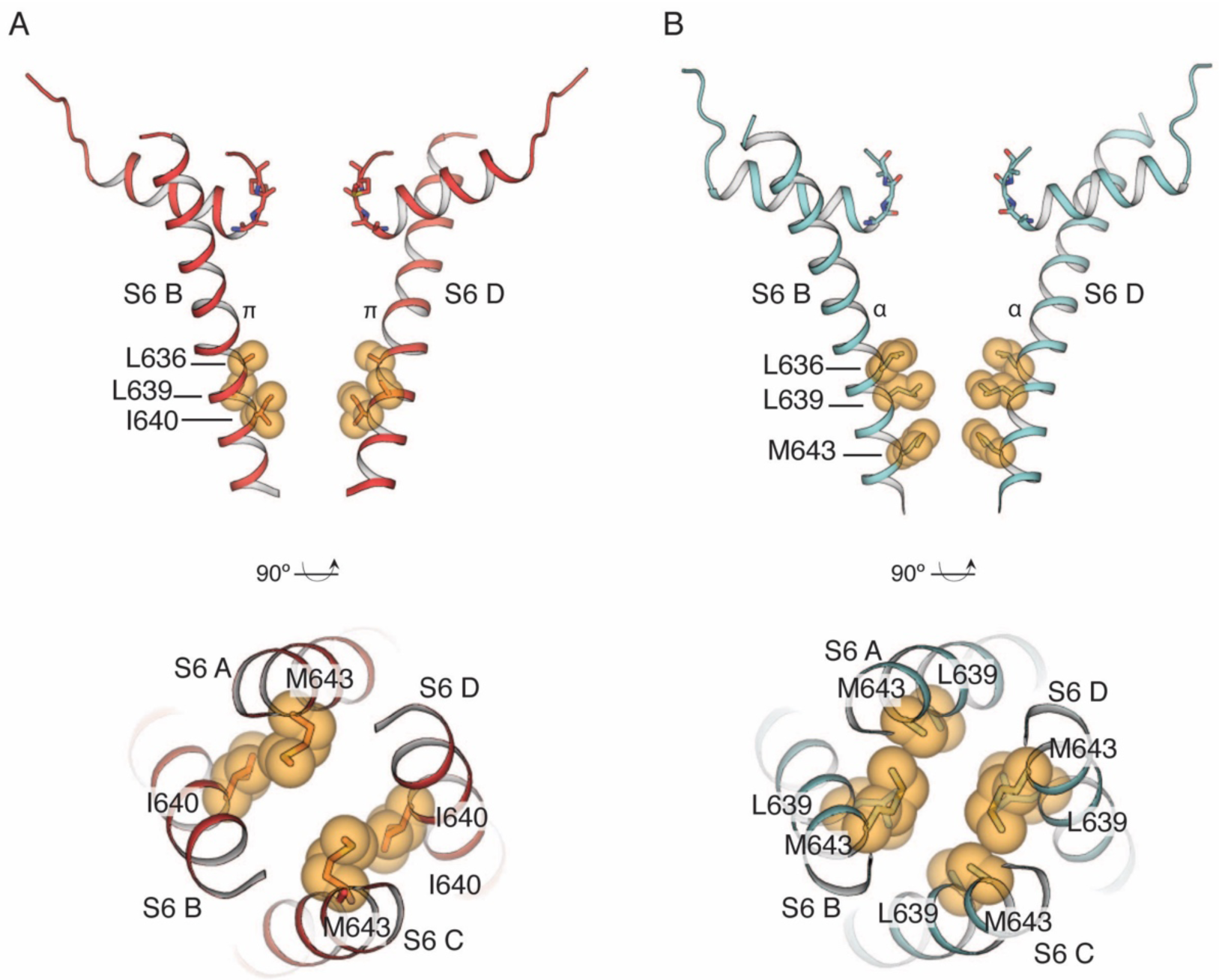
The common gate in TRPV2_RTx-ND_ (red) and TRPV2_RTx-XTAL_ (cyan). **a,** Side view of the TRPV2_RTx-ND_ pore showing subunits B and D (top). Gate residue I640 is shown in yellow spheres, along with the hydrophobic residues L636 and L639 (side chains not built). Bottom-up view (bottom) shows the contribution of all four subunits to the common gate (M643 in subunits A and C, I640 in subunits B and D). **b,** Side view of the TRPV2_RTx-XTAL_ pore, showing subunits B and D (top). Gate residue M643 is shown in yellow spheres, along with hydrophobic pore lining residues L636 and L639. Bottom-up view of the common gate (bottom) shows gate residues M643 (side chain not built in subunits A and C).

## References

1 Clapham, D. E., Runnels, L. W. & Strubing, C. The TRP ion channel family. Nat Rev Neurosci 2, 387–396 (2001).

2 Cao, E., Cordero-Morales, J. F., Liu, B., Qin, F. & Julius, D. TRPV1 channels are intrinsically heat sensitive and negatively regulated by phosphoinositide lipids. Neuron 77, 667–679, doi:10.1016/j.neuron.2012.12.016 (2013).

3 Liu, B. & Qin, F. Use Dependence of Heat Sensitivity of Vanilloid Receptor TRPV2. Biophys J 110, 1523–1537, doi:10.1016/j.bpj.2016.03.005 (2016).

4 Smith, G. D., Gunthorpe, M. J., Kelsell, R. E., Hayes, P. D., Reilly, P., Facer, P., Wright, J. E., Jerman, J. C., Walhin, J. P., Ooi, L., Egerton, J., Charles, K. J., Smart, D., Randall, A. D., Anand, P. & Davis, J. B. TRPV3 is a temperature-sensitive vanilloid receptor-like protein. Nature 418, 186–190, doi:10.1038/nature00894 (2002).

5 Chung, M. K., Lee, H. & Caterina, M. J. Warm temperatures activate TRPV4 in mouse 308 keratinocytes. J Biol Chem 278, 32037–32046, doi:10.1074/jbc.M303251200 (2003).

6 Bolcskei, K., Tekus, V., Dezsi, L., Szolcsanyi, J. & Petho, G. Antinociceptive desensitizing actions of TRPV1 receptor agonists capsaicin, resiniferatoxin and N-oleoyldopamine as measured by determination of the noxious heat and cold thresholds in the rat. Eur J Pain 14, 480–486, doi:10.1016/j.ejpain.2009.08.005 (2010).

7 Julius, D. TRP channels and pain. Annu Rev Cell Dev Biol 29, 355–384, doi:10.1146/annurev-cellbio-101011-155833 (2013).

8 Marwaha, L., Bansal, Y., Singh, R., Saroj, P., Bhandari, R. & Kuhad, A. TRP channels: potential drug target for neuropathic pain. Inflammopharmacology 24, 305–317, doi:10.1007/s10787-016-0288-x (2016).

9 Mitchell, K., Lebovitz, E. E., Keller, J. M., Mannes, A. J., Nemenov, M. I. & Iadarola, M. J. Nociception and inflammatory hyperalgesia evaluated in rodents using infrared laser stimulation after Trpv1 gene knockout or resiniferatoxin lesion. Pain 155, 733–745, doi:10.1016/j.pain.2014.01.007 (2014).

10 Katanosaka, Y., Iwasaki, K., Ujihara, Y., Takatsu, S., Nishitsuji, K., Kanagawa, M., Sudo, A., Toda, T., Katanosaka, K., Mohri, S. & Naruse, K. TRPV2 is critical for the maintenance of cardiac structure and function in mice. Nat Commun 5, 3932, doi:10.1038/ncomms4932 (2014).

11 Eytan, O., Fuchs-Telem, D., Mevorach, B., Indelman, M., Bergman, R., Sarig, O., Goldberg, I., Adir, N. & Sprecher, E. Olmsted syndrome caused by a homozygous recessive mutation in TRPV3. J Invest Dermatol 134, 1752–1754, doi:10.1038/jid.2014.37 (2014).

12 Imura, K., Yoshioka, T., Hirasawa, T. & Sakata, T. Role of TRPV3 in immune response to development of dermatitis. J Inflamm (Lond) 6, 17, doi:10.1186/1476-9255-6-17 (2009).

13 Kim, H. O., Cho, Y. S., Park, S. Y., Kwak, I. S., Choi, M. G., Chung, B. Y., Park, C. W. & Lee, J. Y. Increased activity of TRPV3 in keratinocytes in hypertrophic burn scars with postburn pruritus. Wound Repair Regen 24, 841–850, doi:10.1111/wrr.12469 (2016).

14 Asakawa, M., Yoshioka, T., Matsutani, T., Hikita, I., Suzuki, M., Oshima, I., Tsukahara, K., Arimura, A., Horikawa, T., Hirasawa, T. & Sakata, T. Association of a mutation in TRPV3 with defective hair growth in rodents. J Invest Dermatol 126, 2664–2672, doi:10.1038/sj.jid.5700468 (2006).

15 Imura, K., Yoshioka, T., Hikita, I., Tsukahara, K., Hirasawa, T., Higashino, K., Gahara, Y., Arimura, A. & Sakata, T. Influence of TRPV3 mutation on hair growth cycle in mice. Biochem Biophys Res Commun 363, 479–483, doi:10.1016/j.bbrc.2007.08.170 (2007).

16 Xiao, R., Tian, J., Tang, J. & Zhu, M. X. The TRPV3 mutation associated with the hairless phenotype in rodents is constitutively active. Cell Calcium 43, 334–343, doi:10.1016/j.ceca.2007.06.004 (2008).

17 Masuyama, R., Vriens, J., Voets, T., Karashima, Y., Owsianik, G., Vennekens, R., Lieben, L., Torrekens, S., Moermans, K., Vanden Bosch, A., Bouillon, R., Nilius, B. & Carmeliet, G. TRPV4-mediated calcium influx regulates terminal differentiation of osteoclasts. Cell Metab 8, 257–265, doi:10.1016/j.cmet.2008.08.002 (2008).

18 Chung, M. K., Guler, A. D. & Caterina, M. J. TRPV1 shows dynamic ionic selectivity during agonist stimulation. Nat Neurosci 11, 555–564, doi:10.1038/nn.2102 (2008).

19 Puopolo, M., Binshtok, A. M., Yao, G. L., Oh, S. B., Woolf, C. J. & Bean, B. P. Permeation and block of TRPV1 channels by the cationic lidocaine derivative QX-314. J Neurophysiol 109, 1704–1712, doi:10.1152/jn.00012.2013 (2013).

20 Liao, M., Cao, E., Julius, D. & Cheng, Y. Structure of the TRPV1 ion channel determined by electron cryo-microscopy. Nature 504, 107–112, doi:10.1038/nature12822 (2013).

21 Zubcevic, L., Herzik, M. A., Jr., Chung, B. C., Liu, Z., Lander, G. C. & Lee, S. Y. Cryo-electron microscopy structure of the TRPV2 ion channel. Nat Struct Mol Biol, doi:10.1038/nsmb.3159 (2016).

22 Huynh, K. W., Cohen, M. R., Jiang, J., Samanta, A., Lodowski, D. T., Zhou, Z. H. & Moiseenkova-Bell, V. Y. Structure of the full-length TRPV2 channel by cryo-EM. Nat Commun 7, 11130, doi:10.1038/ncomms11130 (2016).

23 Zhang, F., Hanson, S. M., Jara-Oseguera, A., Krepkiy, D., Bae, C., Pearce, L. V., Blumberg, P. M., Newstead, S. & Swartz, K. J. Engineering vanilloid-sensitivity into the rat TRPV2 channel. Elife 5, doi:10.7554/eLife.16409 (2016).

24 Zubcevic, L., Le, S., Yang, H. & Lee, S. Y. Conformational Plasticity in the Selectivity Filter of the TRPV2 Ion Channel. Nat Struct Mol Biol, in press, DOI:10.1038/s41594-41018-40059-z (2018).

25 Yao, J., Liu, B. & Qin, F. Pore turret of thermal TRP channels is not essential for temperature sensing. Proc Natl Acad Sci U S A 107, E125; author reply E126-127, doi:10.1073/pnas.1008272107 (2010).

26 Jara-Oseguera, A., Bae, C. & Swartz, K. J. An external sodium ion binding site controls allosteric gating in TRPV1 channels. Elife 5, doi:10.7554/eLife.13356 (2016).

27 Dosey, T. L., Wang, Z., Fan, G., Zhang, Z., Serysheva, II, Chiu, W. & Wensel, T. G. Structures of TRPV2 in distinct conformations provide insight into role of the pore turret. Nat Struct Mol Biol 26, 40–49, doi:10.1038/s41594-018-0168-8 (2019).

28 Zoonens, M. & Popot, J. L. Amphipols for Each Season. J Membrane Biol 247, 759–796, doi:10.1007/s00232-014-9666-8 (2014).

29 Cao, E., Liao, M., Cheng, Y. & Julius, D. TRPV1 structures in distinct conformations reveal activation mechanisms. Nature 504, 113–118, doi:10.1038/nature12823 (2013).

30 Paulsen, C. E., Armache, J. P., Gao, Y., Cheng, Y. & Julius, D. Structure of the TRPA1 ion channel suggests regulatory mechanisms. Nature 520, 511–517, doi:10.1038/nature14367 (2015).

31 Yoo, J., Wu, M., Yin, Y., Herzik, M. A., Jr., Lander, G. C. & Lee, S. Y. Cryo-EM structure of a mitochondrial calcium uniporter. Science 361, 506–511, doi:10.1126/science.aar4056 (2018).

32 Hirschi, M., Herzik, M. A., Jr., Wie, J., Suo, Y., Borschel, W. F., Ren, D., Lander, G. C. & Lee, S. Y. Cryo-electron microscopy structure of the lysosomal calcium-permeable channel TRPML3. Nature 550, 411–414, doi:10.1038/nature24055 (2017).

33 Zubcevic, L., Herzik, M. A., Jr., Wu, M., Borschel, W. F., Hirschi, M., Song, A. S., Lander, G. C. & Lee, S. Y. Conformational ensemble of the human TRPV3 ion channel. Nat Commun 9, 4773, doi:10.1038/s41467-018-07117-w (2018).

34 Denisov, I. G. & Sligar, S. G. Nanodiscs for structural and functional studies of membrane proteins. Nat Struct Mol Biol 23, 481–486, doi:10.1038/nsmb.3195 (2016).

35 Scheres, S. H. RELION: implementation of a Bayesian approach to cryo-EM structure determination. J Struct Biol 180, 519–530, doi:10.1016/j.jsb.2012.09.006 (2012).

36 Matthies, D., Dalmas, O., Borgnia, M. J., Dominik, P. K., Merk, A., Rao, P., Reddy, B. G., Islam, S., Bartesaghi, A., Perozo, E. & Subramaniam, S. Cryo-EM Structures of the Magnesium Channel CorA Reveal Symmetry Break upon Gating. Cell 164, 747–756, doi:10.1016/j.cell.2015.12.055 (2016).

37 Yin, Y., Wu, M., Hsu, A., Borschel W.F., Borgnia, M., Lander, G.C., Lee, S-Y. Visualizing structural transitions of ligand-dependent gating of the TRPM2 channel. BioRxiv, doi:https://doi.org/10.1101/516468 (2018).

38 Tonngu, L., Wang, L. Broken symmetry in the human BK channel. BioRxiv, doi:https://doi.org/10.1101/494385 (2018).

39 She, J., Guo, J., Chen, Q., Zeng, W., Jiang, Y. & Bai, X. C. Structural insights into the voltage and phospholipid activation of the mammalian TPC1 channel. Nature 556, 130–134, doi:10.1038/nature26139 (2018).

40 Shen, H., Zhou, Q., Pan, X., Li, Z., Wu, J. & Yan, N. Structure of a eukaryotic voltage-gated sodium channel at near-atomic resolution. Science 355, doi:10.1126/science.aal4326 (2017).

41 Pan, X., Li, Z., Zhou, Q., Shen, H., Wu, K., Huang, X., Chen, J., Zhang, J., Zhu, X., Lei, J., Xiong, W., Gong, H., Xiao, B. & Yan, N. Structure of the human voltage-gated sodium channel Na_V_1.4 in complex with beta1. Science 362, doi:10.1126/science.aau2486 (2018).

42 Shen, H., Li, Z., Jiang, Y., Pan, X., Wu, J., Cristofori-Armstrong, B., Smith, J. J., Chin, Y. K. Y., Lei, J., Zhou, Q., King, G. F. & Yan, N. Structural basis for the modulation of voltage-gated sodium channels by animal toxins. Science 362, doi:10.1126/science.aau2596 (2018).

43 Hille, B. The permeability of the sodium channel to organic cations in myelinated nerve. J Gen Physiol 58, 599–619 (1971).

44 Ritchie, T. K., Grinkova, Y. V., Bayburt, T. H., Denisov, I. G., Zolnerciks, J. K., Atkins, W. M. & Sligar, S. G. Chapter 11 - Reconstitution of membrane proteins in phospholipid bilayer nanodiscs. Methods Enzymol 464, 211–231, doi:10.1016/S0076-6879(09)64011-8 (2009).

45 Zheng, S. Q., Palovcak, E., Armache, J. P., Verba, K. A., Cheng, Y. & Agard, D. A. MotionCor2: anisotropic correction of beam-induced motion for improved cryo-electron microscopy. Nat Methods 14, 331–332, doi:10.1038/nmeth.4193 (2017).

46 Zhang, K. Gctf: Real-time CTF determination and correction. Journal of structural biology 193, 1–12, doi:10.1016/j.jsb.2015.11.003 (2016).

47 Zivanov, J., Nakane, T., Forsberg, B. O., Kimanius, D., Hagen, W. J. H., Lindahl, E. & Scheres, S. H. W. New tools for automated high-resolution cryo-EM structure determination in RELION-3. Elife 7, doi:ARTN e4216610.7554/eLife.42166 (2018).

48 Scheres, S. H. & Chen, S. Prevention of overfitting in cryo-EM structure determination. Nat Methods 9, 853–854, doi:10.1038/nmeth.2115 (2012).

49 Chen, S., McMullan, G., Faruqi, A. R., Murshudov, G. N., Short, J. M., Scheres, S. H. & Henderson, R. High-resolution noise substitution to measure overfitting and validate resolution in 3D structure determination by single particle electron cryomicroscopy. Ultramicroscopy 135, 24–35, doi:10.1016/j.ultramic.2013.06.004 (2013).

50 Emsley, P. & Cowtan, K. Coot: model-building tools for molecular graphics. Acta Crystallogr D Biol Crystallogr 60, 2126–2132, doi:S0907444904019158 [pii]10.1107/S0907444904019158 (2004).

51 Adams, P. D., Afonine, P. V., Bunkoczi, G., Chen, V. B., Davis, I. W., Echols, N., Headd, J. J., Hung, L. W., Kapral, G. J., Grosse-Kunstleve, R. W., McCoy, A. J., Moriarty, N. W., Oeffner, R., Read, R. J., Richardson, D. C., Richardson, J. S., Terwilliger, T. C. & Zwart, P. H. PHENIX: a comprehensive Python-based system for macromolecular structure solution. Acta Crystallogr D Biol Crystallogr 66, 213–221, doi:S0907444909052925 [pii]10.1107/S0907444909052925 (2010).

52 Chen, V. B., Arendall, W. B., 3rd, Headd, J. J., Keedy, D. A., Immormino, R. M., Kapral, G. J., Murray, L. W., Richardson, J. S. & Richardson, D. C. MolProbity: all-atom structure validation for macromolecular crystallography. Acta Crystallogr D Biol Crystallogr 66, 12–21, doi:10.1107/S0907444909042073 (2010).

53 Smart, O. S., Neduvelil, J. G., Wang, X., Wallace, B. A. & Sansom, M. S. HOLE: a program for the analysis of the pore dimensions of ion channel structural models. J Mol Graph 14, 354–360, 376 (1996).

54 Pettersen, E. F., Goddard, T. D., Huang, C. C., Couch, G. S., Greenblatt, D. M., Meng, E. C. & Ferrin, T. E. UCSF chimera - A visualization system for exploratory research and analysis. J Comput Chem 25, 1605–1612, doi:10.1002/jcc.20084 (2004).

